# CD4^+^ T cell fate decisions are stochastic, precede cell division, depend on GITR co-stimulation, and are associated with uropodium development

**DOI:** 10.1101/298026

**Authors:** Stephen P. Cobbold, Elizabeth Adams, Duncan Howie, Herman Waldmann

## Abstract

During an immune response, naïve CD4^+^ T cells proliferate and generate a range of effector, memory and regulatory T cell subsets, but how these processes are co-ordinated remains unclear. A traditional model suggests that memory cells use mitochondrial respiration and are survivors from a pool of previously proliferating and glycolytic, but short-lived effector cells. A more recent model proposes a binary commitment to either a memory or effector cell lineage during a first, asymmetric cell division, with each lineage able to undergo subsequent proliferation and differentiation. We used improved fixation and staining methods with imaging flow cytometry in an optimised in vitro system that indicates a third model. We found that cell fates result from stochastic decisions that depend on GITR co-stimulation and which take place before any cell division. Effector cell commitment is associated with mTORC2 signalling leading to uropodium development, while developing memory cells lose mitochondria, have a nuclear localization of NFκB and depend on TGFβ for their survival. Induced, T helper subsets and foxp3^+^ regulatory T cells were found in both the effector and memory cell lineages. This in vitro model of T cell differentiation is well suited to testing how manipulation of cytokine, nutrient other components of the microenvironment might be exploited for therapeutic purposes.

## 1. Introduction

A fundamental feature of the adaptive immune system is the ability to mount a rapid and protective, secondary or memory response to a pathogen it has previously encountered. This memory derives from both an increase in the frequency of pathogen specific lymphocytes by clonal expansion together with their differentiation into long-lived memory cells that can rapidly generate the most appropriate secondary effector functions. As there is no affinity maturation in T cells, antigen specific T cells must be able to generate both the short-term, terminally differentiated effector cells for the primary response as well as long-lived, protective memory cells. The mechanisms by which these two distinct populations are generated from homogenous clones of naïve T cells remain poorly understood.

There are two prevalent hypotheses in the literature to explain how naïve T cells generate both short-term effector and long-term memory cells. The first is a linear model where activated T cells first proliferate, driven by cytokines such as IL-2, mTOR activation and glycolysis, to generate a large population of effector cells (1, 2). Once antigen is cleared, these effectors mostly die leaving a smaller population of surviving T cells that, in the presence of cytokines such as IL-15, fatty acid driven oxidative phosphorylation (3), and TNFRSF (CD27, CD134) signalling (4, 5) further differentiate into long-term memory T cells. Recently, a second, more controversial (6, 7) model suggests that after activation, T cells undergo one or more polarised or asymmetric cell divisions, with one daughter destined to become a short-lived effector cell while the other develops into a long-lived memory cell (8, 9). These alternate fates are said to be determined by an asymmetric inheritance during cytokinesis of the numb/notch signalling pathway (10), cell surface molecules such as CD4 and CD8 (9, 11), transcription factors such as Tbet (8), and nutrient sensing pathways via PI3k and CD98 (12-14), which then contribute to the metabolic programming towards glycolysis in effector cells (15) and to oxidative phosphorylation in memory cells. Most of the published evidence in favour of either model concerns naïve CD8^+^ T cells differentiating into cytotoxic effector versus memory T cells, but similar claims of asymmetric cell divisions are also emerging for conventional CD4^+^ T cells (16).

Peripheral antigen specific, foxp3^+^ expressing CD4^+^ Treg cells are known to result from stimulation of naïve CD4^+^ T cells in the presence of TGFβ, acting via a response element in CNS1 of the foxp3 locus (17). Very little is known about such induced Treg cells in the context of effector versus memory cell fate decisions (18), perhaps because the literature has concentrated on Treg cell development and repertoire selection in the thymus (19, 20). Peripherally induced, antigen specific CD4^+^foxp3^+^ Treg cells are important for immune regulation and are required for certain forms of transplantation tolerance (21). Tolerance is not simply that T cell development has switched from an effector to regulatory cell fate, as tolerant mice can still sustain a large population of effector cells (22). In these circumstances, regulatory T cells and conventional memory cells have both previously been exposed to their antigen and both seem to depend on fatty acids and oxidative phosphorylation when compared to activated naïve cells and effector cells (23, 24). This might indicate some commonality in the mechanisms that determine the memory T cell and Treg cell fates. By analogy with models for conventional effector versus memory cell fate decision, peripheral Treg cells could also develop either as survivors from a previously proliferating effector cell population (25) or from a binary cell fate decision during an asymmetric cell division (9).

To study CD4^+^ T cell fate decisions, we needed a system where we could control multiple stimuli through the TCR, co-stimulation, cytokines and nutrient availability, which together signal through overlapping and non-linear pathways. We did not want to restrict our observations to a binary outcome as there is the potential to generate a range of different effector or memory cell populations with different probabilities. We developed an in vitro culture system to simultaneously study a range of cell fates such as proliferation versus cell death, effector versus memory commitment and conventional versus regulatory T cell subset differentiation, so that we could relate these to TCR, costimulatory, cytokine and nutrient sensing signalling pathways, at both the single cell and population levels. To achieve this we optimised multicolour staining methods together with imaging flow cytometry (26) so as to track and quantitate the complex outcomes from antigen driven stimulation in vitro of an uniform population of monoclonal, naïve CD4^+^ T cells, and where we could control and manipulate the culture and stimulation conditions.

We used imaging flow cytometry as it is ideally suited to simultaneously quantifying multiple parameters at both the single cell and population level so allowing us to combine staining for markers of cell differentiation with structural information, such as the shape or asymmetry of cells, together with the localisation and polarisation of cell surface and nuclear markers, and of intracellular organelles such as mitochondria. The role of cell structure in lymphocyte function has been little studied at the population level, with conventional flow cytometry indicating only the size (forward scatter) and complexity (side scatter). One of the most microscopically obvious structures is the uropodium (27, 28), which is a large protrusion at the rear of lymphocytes migrating on an appropriate matrix. As well as a role in migration, it has been proposed that uropodia are important in interactions with antigen presenting cells, cytotoxicity, and cell fate decisions (28). Uropodia are dynamic structures requiring active maintenance of the cytoskeleton and microtubules, essential for effective immune responses in vivo, yet their role in lymphocytes is still poorly understood. Uropodia contain the bulk of the cytoplasm and organelles such as the microtubule organising centre (MTOC), mitochondria (29), lysosomes and golgi. Many of the cell surface molecules involved in interactions with other cells, including components of the immune synapse and CD44 are also localised to the uropodium (28).

We use a multi-dimensional analysis of many individual CD4^+^ T cells in terms of their proliferation history, differentiation, cell structure, signalling and survival in response to the microenvironment within which they are stimulated. The data we presentsupport a model that favours initial, stochastic cell fate commitments, for both conventional and regulatory CD4^+^ T cells, that are dependent on multiple interacting signalling pathways during their initial activation. These act to determine the cell fate before the commencement of cell division, which then takes place entirely symmetrically to generate two identically committed daughters. Both the effector and memory cell populations proliferate. Effector cells die after 4-5 cell divisions, while memory cell survive and enter quiescence as mTOR signalling decays as antigen is cleared or nutrients, such as amino acids, become limiting (30). These memory cells can then make further fate decisions upon secondary stimulation.

## 2. Materials and Methods

### 2.2 Mice

A1.RAG1^−/−^ (TCR transgenic anti-Dby+IE^k^; on a CBA/Ca.RAG1^−/−^ background:”A1RAG”) (31), CBA/Ca, CBA.RAG1^−/−^, Marilyn.hCD2-Foxp3.RAG1^−/−^ (TCR transgenic anti-Dby+IA^b^ on a C57BL/6,RAG1^−/−^ background with hCD2-Foxp3 reporter: “MARKI”) (21) and C57BL/6J mice were bred and maintained under SPF conditions in the animal facility of the Sir William Dunn School of Pathology, Oxford, UK. All procedures were conducted in accordance with the Home Office Animals (Scientific Procedures) Act of 1986 (PPL 30/3060).

### 2.2 Skin grafting

A1RAG female mice of 6-8 weeks of age were given male CBA.RAG1^−/−^ skin grafts (31). Control mice were allowed to reject these grafts while the tolerant (31) group received 1mg of YTS 177 on day 0 and maintained their grafts until the end of the experiment. All mice were given second challenge male CBA.RAG1^−/−^ skin grafts after 3 months and were sacrificed 7 days later and their draining lymph nodes were taken and prepared for staining with Mitotracker DR and antibody markers (Table 1).

**Table 1:** Fluorescent reagents for Imaging Flow Cytometry

| ISX | Stain Type | Reagents used |
| --- | --- | --- |
| *Ch1* | *N/A* | *Bright field camera 1* |
| Ch2 | Fix/Perm | AlexaFluor488 Mouse anti-Tbet (BD Pharmingen 561266) |
| **Ch2** | **Fix/Perm** | **Rabbit mAb anti-phospho-S6-AlexaFluor488 (Cell Signaling Technology #4854)** |
| Ch2 | Fix/Perm | Rabbit mAb anti-pan AKT-AlexaFluor488 (Cell Signaling Technology #5084S) |
| Ch2 | Fix/Perm | Rabbit mAb anti-pAKT<sub>T308</sub>-AlexaFluor488 (Cell Signaling Technology #2918S) |
| Ch2 | Fix/Perm | Mouse anti-gamma tubulin-AlexaFluor488 (clone TU-30; EXBIO A4-465-C100) |
| Ch2 | Brefeldin/Perm | Rat anti-mouse IL2-FITC (BD Pharmingen 554427) |
| Ch2 | Live cell | Rat anti-mouse CD11a/LFA-1-FITC (eBioscience 11-0111-85) |
| Ch2 | Fix/Perm | AlexaFluor488 Rat anti-mouse GITR (BioLegend 120211) |
| Ch2 | Fix/perm | FITC anti-mouse Fas (BD Pharmingen 15404D) |
| Ch2 | Fix/Perm | SYTOX Green (DNA) (Life Technologies S7020) |
| Ch2 | Live/Fix/Acquire | Autophagy Green Detection Reagent (Abcam 139484) |
| **Ch3** | **Fix/Perm** | **Rabbit mAb anti-pAKT<sub>S473</sub>-PE (Cell Signaling Technology #5315S)** |
| Ch3 | Fix/Perm | Anti-human/mouse GATA3-PE (eBioscience 12-9966-42) |
| Ch3 | Fix/Perm | Anti-human/mouse IRF4-PE (eBioScience 12-9858-82) |
| Ch3 | Fix/Perm | Anti-mouse granzyme B-PE (eBioscience 12-8898-82) |
| Ch3 | Brefeldin/Perm | Rat anti-mouse IFNγ-PE (BD Pharmingen 554412) |
| Ch3 | Live cell | PE anti-mouse/rat CD29 (BioLegend 102208) |
| Ch3 | Live cell | PE anti-mouse FasL (eBioscience 12-5911-82) |
| Ch3 | Fix/Perm | Mouse anti-human Ki67-PE (BD Pharmingen 51-36525X) |
| **Ch4** | **Live or fix/perm** | **PE-CF594 rat anti-mouse CD4 (BD Horizon 562314)** |
| Ch4 | Fix/Perm | PE-CF594 anti-RORγt (BD Horizon 562884) |
| Ch4 | Brefeldin/Perm | PE-CF594 anti-IL4 (BD Horizon 562450) |
| **Ch5** | **Fix/Perm** | **7AAD (DNA) (Sigma A9400-5MG)<sup>a</sup>** |
| Ch5 | Fix/Perm | Mito-ID-Red (Enzo ENZ-51007-500)<sup>a</sup> |
| **Ch6** | **Fix/Perm** | **PE/Cy7 rat anti-mouse Foxp3 (eBioscience 25-5773-82)** |
| Ch6 | Live cell | PE/Cy7 anti-CD27 (eBioscience 25-0271-82) |
| **Ch7** | **Live cell** | **Cell Trace Violet (CTV: Invitrogen C34557)** |
| **Ch8** | **Live cell** | **Fixable LIVE/DEAD Aqua (Life technologies L34957)** |
| *Ch9* | *N/A* | *Bright field camera 2* |
| **Ch10** | **Live cell** | **Rat anti-mouse CD62L Brilliant Violet 605 [“BV605”] (BioLegend 104437)** |
| Ch10 | Live cell | Anti-mouse CD25 Brilliant Violet 605 (BioLegend 102036) |
| Ch10 | Fix/Perm | Rat anti-mouse Tbet Brilliant Violet 605 (BioLegend 644817) |
| Ch10 | Brefeldin/Perm | Rat anti-mouse IL17 Brilliant Violet 605 (BioLegend 506927) |
| **Ch11** | **Live cell** | **Mitotracker Deep Red FM (MitoDR: Invitrogen M22426)** |
| **Ch11** | **Fix/Perm** | **Anti-NFkB p65 AlexaFluor 647 (Abcam ab 190589)** |
| **Ch12** | **Live cell** | **Anti-human/mouse CD44 APC-eFluor780 (eBioscience 47-0441-82)** |
| Ch12 | Live cell | Anti-mouse CD4 APC-Cy7 (Biolegend 100526) |
| Ch12 | Fix/perm | Anti-mouse CD3ε APC-Cy7 (BD Pharmingen 557596) |
| Ch12 | Live cell | Anti-mouse CD25 APC-eFluor780 (eBioscience 47-0251-82) |

Lasers powers: 405nm (50mW), 488nm (100mW), 560nm (**OFF**), 642nm (100mW), SSC (**OFF**)
<sup>a</sup> 7AAD or Mito-Id-Red must be excited by 488nm to give weak emission in Ch4 and Ch5, so that compensation can be achieved with strong emission of PE-CF594 in Ch4.
**Reagents in bold show the most frequently used combination for 10 colour staining panel.**

### 2.3 TCR transgenic T cells and Treg cell cultures

CD4^+^ TCR transgenic T cells were selected from spleen cells of female A1RAG or MARKI mice using the CD4 isolation kit (Miltenyi Biotec: 130-104-454) and were labelled with Cell Trace Violet (CTV: Invitrogen C34557)), according to the manufacturers’ instructions. These labelled CD4^+^ T cells were cultured at 5x10^5^ cells/ml in 48x1ml tissue culture plates in Advanced RPMI 1640 (Life Technologies 12633-020) with added GlutaMAX (Life technologies 35050), 10^−5^ M mercaptoethanol, 10mM HEPES, a reduced (1/10) concentration of penicillin/streptomycin, plus 1% FCS together with either 1x10^5^ syngeneic bmDC (32, 33) plus the appropriate Dby peptide (Dby-E^k^: REEALHQFRSGRKPI: 100nM unless otherwise stated, Dby-A^b^: NAGFNSNRANSSRSS: 10nM) (21, 31) or with CD3/CD28 beads (Dynabeads Mouse T-activator CD3/CD28: Life Technologies 11452D) at 1:1 ratio or with anti-CD3 (145-2C11: 0.1-5μg/ml or as stated, coated on plastic in 0.1M sodium bicarbonate) plus soluble anti-CD28 (1μg/ml clone 37.51) for 3 days (A1RAG cells) or 2 days (MARKI cells), unless time point stated otherwise, in a gassed, humidified incubator at 37°C plus 5% CO_2_. All cultures included 50U/ml rmIL-2, and 2ng/ml rhTGFβ (Peprotech 100-21C) plus 100nM all trans-retinoic acid (ATRA: Sigma R2625), unless stated otherwise. Inhibitors were added where indicated: rapamycin (Calbiochem 553211: 50nM), Torin1 (Tocris Bioscience 4247: 250nM), anti-LFA-1 (FD441.8; 50μg/ml) (34), anti-GITR [YGITR 765.4 (35, 36) 50μg/ml], anti-GITRL [YGL 386.2 (35) 50μg/ml].

### 2.4 Staining cells for 10 colour flow cytometric imaging

If the staining included anti-cytokine antibodies then Brefeldin A (5μg/ml) was added for the last 2 hours of cell culture. Staining for fixation sensitive cell surface markers (eg. CD25) and live cell stains were performed in situ, with minimal disturbance of the cells, by adding 100μl of Advanced RPMI 1640 containing 5ng/ml Mitotracker DR (Invitrogen M22426) plus 2μl of live/dead aqua (Life technologies L34957: 1 vial reconstituted in 40μl of DMSO), together with 1μg of each antibody conjugate (Table 1), to each 1 ml of culture, and incubated in the dark, in a humidified gassed (5% CO_2_) incubator at 37°C for 30-60 mins. If samples were to be analysed for mitotic cells in telophase, a pre-warmed (37°C) 40% solution of formaldehyde was added directly to the cultures to a final concentration of 4% formaldehyde and incubated at 37°C for 15 mins. Approx. 95% of the medium was then carefully aspirated without disturbing the cells and 200μl of warm (37°C) fix/permeabilisation buffer for Foxp3 staining (eBioscience 00-5123-43) added and incubated at 37°C in the dark for 2 hours. 1 ml of 1x Foxp3 permeabilisation buffer (eBioscience 00- 8333-56) was then added, the cells were thoroughly re-suspended by vigorous pipetting, harvested and pelleted for labelling with antibody conjugates to fixation resistant cell surface epitopes (eg. CD4, GITR), if required, together with other required antibody conjugates (Table 1) in 1x permeabilisation buffer at room temperature for 1 hour. After washing in 1x permeabilisation buffer, cells were re-suspended in 15μl of PBS + 1% BSA + 0.1% NaN_3_ and fixed by addition of an equal volume of PBS + 4% formalin together with a DNA stain eg. 7- actinomycin D (7AAD: 10μg/ml).

### 2.5 Imaging Flow Cytometry

Samples were run on a 2 camera, 12 channel ImageStream X MkII (Amnis Corporation) with the 60X Multimag objective and the Extended Depth of Field (EDF) option providing a resolution of 0.3μm per pixel and 16μm depth of field. Fluorescent excitation lasers and powers used were 405nm (50mW), 488nm (100mW) and 643nm (100mW) and the side scatter laser was turned off to allow channel 6 to be used for PE-Cy7. The 448nm laser must be used to excite all fluorophores emitting in channels 2-6, as any use of the 560nm laser compromised multicolour compensation. Bright field images were captured on channels 1 and 9 (automatic power setting). A minimum of 30,000 images were acquired per sample using INSPIRE 200 software (Amnis Corporation). Images containing beads were excluded during acquisition as low intensity and high modulation of bright field channels 1 and 9. Images were analysed using the IDEAS v 6.2 software (Amnis Corporation).

### 2.6 Analysis of flow cell images

A colour compensation matrix was generated for all 10 fluorescence channels using samples stained with single colour reagents or antibody-conjugate coated compensation beads, run with the INSPIRE compensation settings, and analysed using the IDEAS compensation wizard. Note that is important to generate a new compensation matrix for each unique combination of fluorophores, and particular caution must be taken with the use of 7AAD or Mito-ID-Red, which emit in both channels 4 and 5 and overlap with PE-CF594. All images were initially gated for focus (using the Gradient RMS feature) on both bright field channels (1 and 9) followed by selecting for singlet cells (DNA intensity/aspect ratio) and live cells at the time of staining ie. live/dead aqua low intensity (channel 8) or low bright field contrast (channel 1).

### 2.7 Identification and measurement of uropodia and associated stains

The strategy for making masks (blue shading) for nuclear expression and identification of uropodia is shown in **Figure 1A**. The uropodium mask relies on the fact that irregular shaped cells tend to align to the direction of laminar flow so the uropodium appears as a protrusion that is aligned to plane of the 2D image of the cell. The bright field (Ch01) default mask (M01) was first eroded, either by 2 pixels or, when a significant number of images contained a lot of extraneous material (as seen at the bottom right of the example shown), by using an 80% adaptive erode (a) followed by a 5 pixel dilation, to generate a “clean” cell mask (b). The nuclear mask was then made using the morphology function of the DNA (eg. 7AAD, Ch05, mask M05, shown in c). The uropodium mask (f) was defined as the largest area single component of the clean cell mask (b) after subtraction of the nuclear mask (c) dilated by 6 pixels (d). **B-G**: Image gating strategy for defining cells with uropodia. Images were gated for focus (**B**), size (bright field area) and non-apoptotic (low bright field contrast) (C). Note that it is essential that images are gated on diploid (ideally G_0_/G_1_ DNA staining intensity) with an aspect ratio >0.8 (ie a single, round nucleus) (**D**). Dead cells staining with live/dead aqua were excluded (**E**). The area of the uropodium mask for each image was then plotted on a frequency histogram (with a log scale for uropodium area) and cells with uropodium areas greater or less than 10μm^2^ were defined as uropodia positive (red) or negative (blue), respectively (**F**). The uropodium mask was also used to calculate the proportion, as a percentage, of any stains of interest (calculated as 100 × intensity of stain within uropodium mask/total stain intensity: an example is shown for mitochondria in **G**).

**Figure 1:**
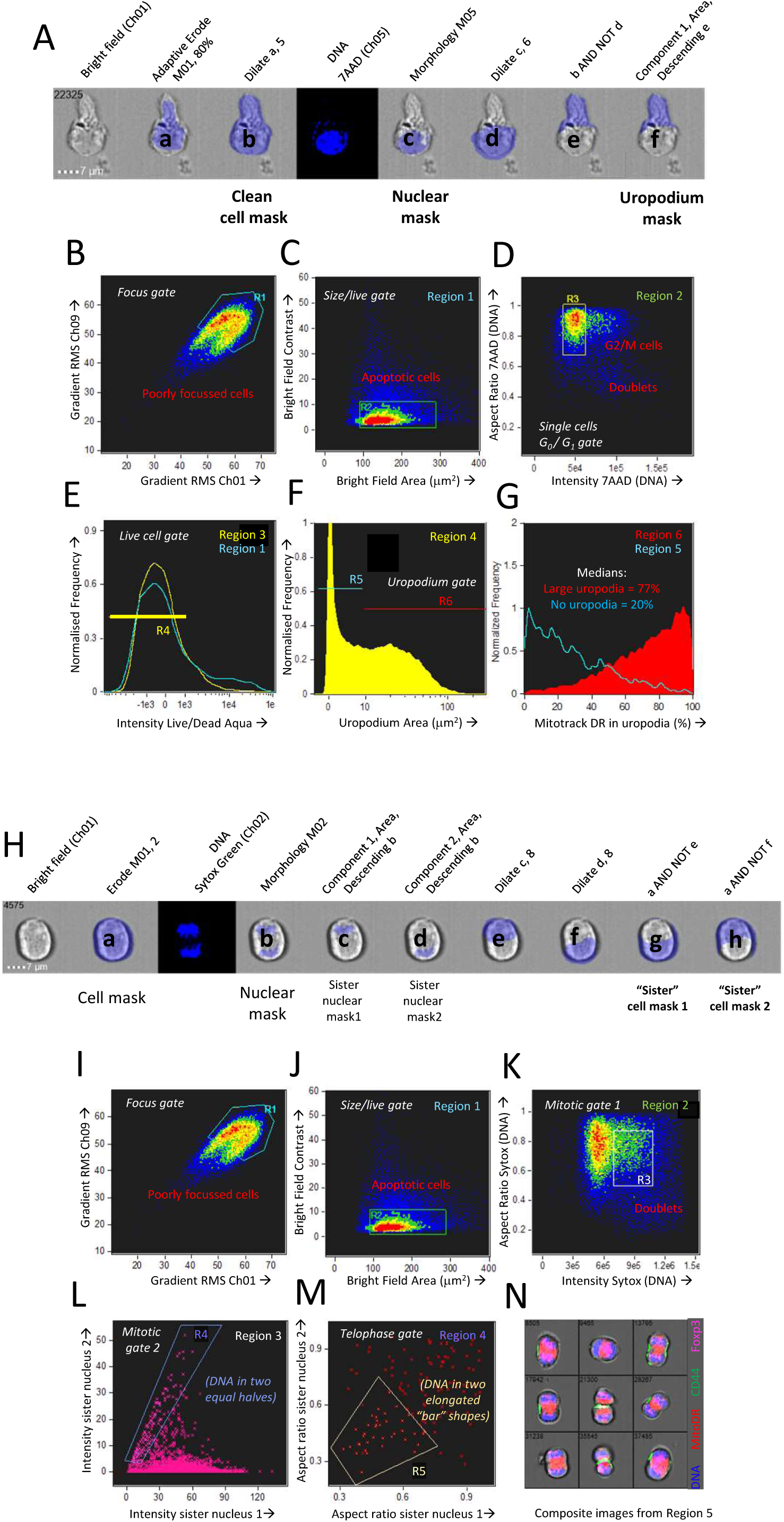
Masking and gating strategies for analysis of uropodia and cells in telophase. The strategy for making masks (blue shading) for nuclear expression and identification of uropodia is shown in **A**. The bright field (Ch01) default mask (M01) was first eroded, either by 2 pixels or, when a significant number of images contained a lot of extraneous material (as seen at the bottom right of the example shown), by using an 80% adaptive erode (a) followed by a 5 or 6 pixel dilation, to generate a “clean” cell mask (b). The nuclear mask was then made using the morphology function of the DNA (7AAD, Ch05, mask M05, shown in c). The uropodium mask (f) was defined as the largest area single component of the clean cell mask (b) after subtraction of the nuclear mask (c) dilated by 6 pixels (d). **B-G**: Image gating strategy for defining cells with uropodia. Images were gated for focus (**B**), size (bright field area) and non-apoptotic (low bright field contrast) (C). Note that it is essential that images are gated on diploid (ideally G_0_/G_1_ DNA staining intensity) with an aspect ratio >0.8 (ie a single, round nuclei) (**D**). Dead cells staining with live/dead aqua were excluded (**E**). The area of the uropodium mask for each image was then plotted on a frequency histogram (with a log scale for uropodium area) and cells with uropodia areas greater or less than 10μm^2^ were defined as uropodia positive (red) or negative (blue), respectively (**F**). The uropodium mask was also used to show the proportion (%) of any stains of interest that were differentially expressed within or outside the uropodium mask (an example is shown for mitochondria in **G**). **H-N**: The strategy for making masks (blue shading) to identify and determine the polarity of telophase cells is shown. A cell mask was made by eroding the default bright field mask (M01) by 2 pixels (a). A nuclear mask was generated by applying the morphology function to the default DNA channel (Ch02, M02 for Sytox Green shown). The component function (Component 1 and 2 sorted for largest area) was then used to identify the DNA staining for the two condensed sister nuclei (c, d) which were dilated by 8 pixels (e, f) and then each was subtracted from the cell mask (a) to give the two sister “cell masks” (g, h). The gating strategy for identify cells in telophase (and late anaphase) is shown in **I-N**. Images were gated for focus (**I**), size (bright field area) and non-apoptotic (low bright field contrast) (**J**), singlet cells with G_2_/M DNA content (**K**) and live cells excluding live/dead aqua (not shown). Cells in late anaphase and telophase were selected by gating for images with two nuclear components of similar DNA stain intensity (**L**) and low aspect ratios (ie. with condensed “bar” shaped nuclei: **M**), with examples shown in **N** (DNA in blue, mitochondria in red and CD4 in green).

### 2.8 Flow imaging analysis of mitotic cells in telophase

We did not use any mitotic inhibitors to enhance the frequency of cells in telophase as these risk inducing asymmetric artefacts in mitotically arrested cells (37). Cell images were gated for focus and live cells as above. The strategy for making masks (blue shading) to identify and determine the polarity of telophase cells is shown in **Figure 1H**. A cell mask was made by eroding the default bright field mask (M01) by 2 pixels (a). A nuclear mask was generated by applying the morphology function to the default DNA channel (eg. Ch02, M02 for Sytox Green shown). The component function (Component 1 and 2 sorted for largest area) was then used to identify the DNA staining for the two condensed sister nuclei (c, d) which were dilated by 8 pixels (e, f) and then each was subtracted from the cell mask (a) to give the two sister “cell masks” (g, h). The gating strategy to identify cells in telophase (and late anaphase) is shown in **Figure 1 I-N**. Images were gated for focus (**I**), size (bright field area) and non-apoptotic (low bright field contrast) (**J**), singlet cells with G^2^/M DNA content (**K**) and live cells excluding live/dead aqua (not shown). Cells in late anaphase and telophase were selected by gating for images with two nuclear components of similar DNA stain intensity (**L**) and low aspect ratios (ie. with condensed “bar” shaped nuclei: **M**), with examples shown in **N** (DNA in blue, mitochondria in red and CD4 in green). A polarity score (with 0 = equal distribution and 100 = all staining within one half) was calculated as:

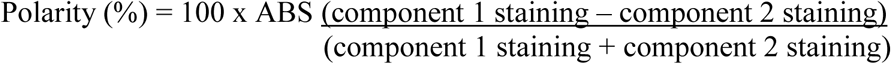

Nuclear intensity was calculated using the intensity feature subject to the nuclear (morphology) mask.

### 2.9 Statistics

Statistical analyses used Prism v 7 (GraphPad) to determine 2 tailed P values by unpaired t test with Welch’s correction to compare 2 groups or ANOVA with Dunnet’s multiple comparison post-test where there were more than 2 groups. Unless otherwise stated, error bars indicate SD. Summary statistics are presented as median values (eg. median fluorescence intensity: MFI), and, where appropriate, a robust CV (%) is indicated.
Mean numbers of cell divisions (Divs) was calculated as:

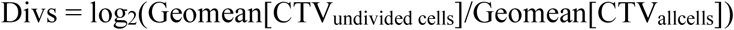

### 2.10 Data availability

Complete original raw image, compensated image, compensation matrix and IDEAS data analysis files for all datasets presented are available from the corresponding author upon request.

## 3. Results

### 3.1 Optimisation of an in vitro system for the activation and differentiation of naïve CD4^+^ T cells

A number of publications have described in vitro cultures for following T cell fate decisions after antigen stimulation, where the effector and memory cells fates can be distinguished by differences in the surface expression of CD4 or CD8, differential PI3k/mTOR signalling, and numbers or activity of mitochondria (9, 13, 38). We used these observations to guide the optimisation of an experimental setup summarised in Figure 2A. TCR transgenic, RAGKO mice provided monoclonal populations of uniformly naïve CD4^+^ T cells that could be analysed in detail after stimulation by their cognate antigen (the male antigen Dby). The A1RAG strain (31) used for most of the in vitro studies reported here, has a low affinity TCR expressed on the CBA/Ca background, but we also reproduced our findings with the Marilyn.hCD2-Foxp3 (MARKI) strain (21) which has a TCR affinity approx. 10 times higher (ie. requires 10 fold less peptide to achieve an equivalent response) and is on the C57BL/6 background.

**Figure 2:**
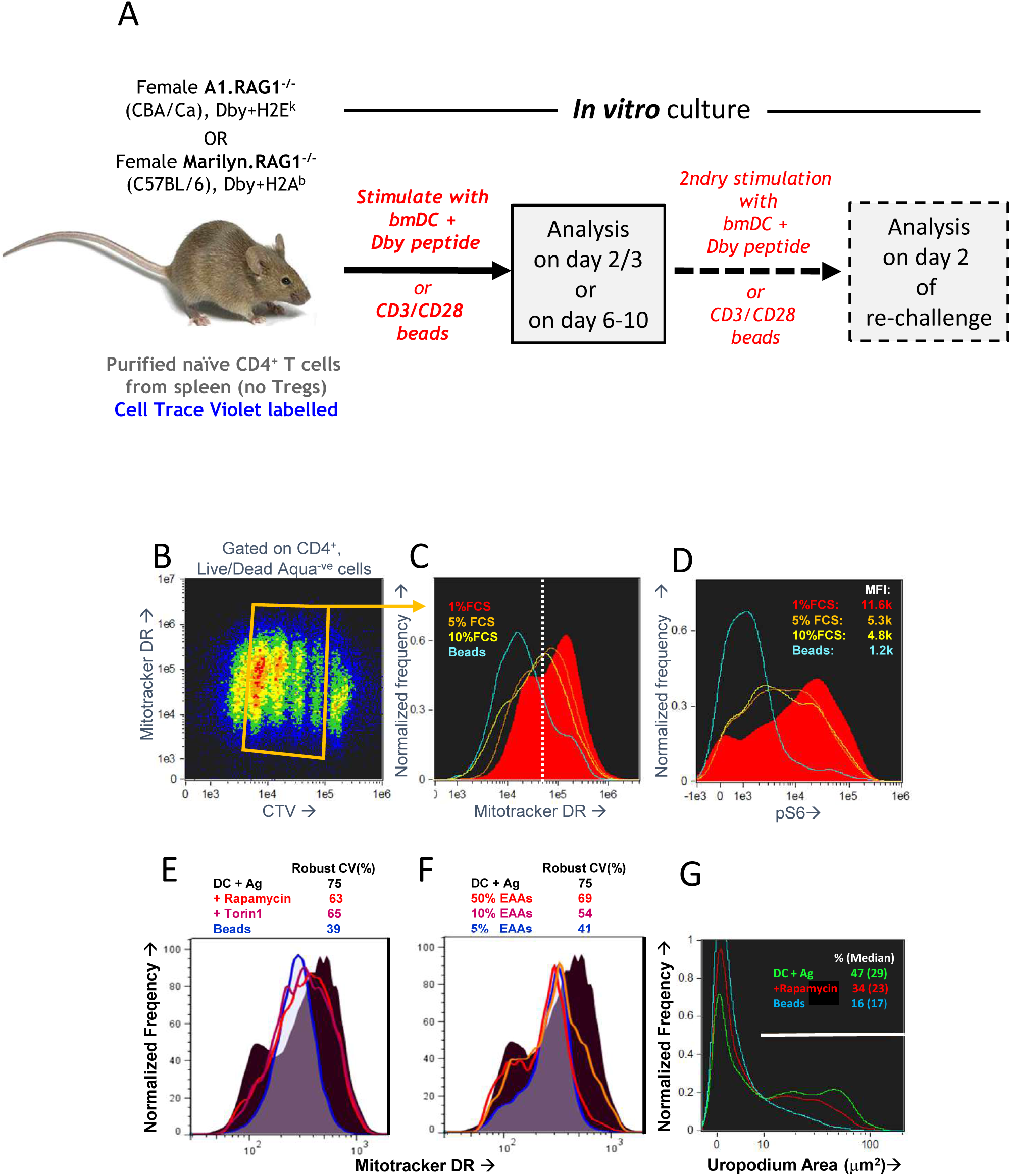
Optimisation of an in vitro culture system of antigen specific stimulation that recapitulates the bimodal distribution of mitochondria observed in vivo. **A**: Schematic of an in vitro system to track the activation, proliferation and differentiation of naïve CD4^+^ T cells at different time points after a primary stimulation, and after a secondary activation. **B-D**: Naïve CD4^+^ T cells from A1RAG mice were labelled with Cell Trace Violet (CTV) and stimulated in vitro with either bmDC plus 100nM Dby peptide or CD3/CD28 beads (blue lines), in the presence of IL-2, TGFβ and ATRA, for 3 days in Advanced RPMI with either 1% (red lines), 5% (orange lines), or 10% FCS (yellow lines) followed by “in situ” staining (see methods) for Mitotracker DR, live/dead Aqua, CD4-PE-CF594 and CD44- APC-eflour780, fixation/permeabilisation, followed by intracellular pS6-Alexa488 and 7AAD staining. ImageStream analysis was performed on 30,000 images per sample, gating on singlet cells as above, CD4+ and live/dead aqua negative, and plotting CTV against Mitotracker DR (**B**: density plot of all samples pooled). Individual samples were gated on cells that had divided 1-4 times (orange box) and the intensity histograms of Mitotracker DR (**C**) and pS6 (**D**) are shown. **E-F**: CTV labelled A1RAG CD4^+^ T cells were stimulated as above with bmDC + Dby peptide, but in standard RPMI + 10% dialysed FCS, or with RPMI with reduced levels of essential amino acids (EAAS), or with addition of mTOR inhibitors rapamycin or Torin 1, as indicated. CD3/CD28 bead stimulation was used as a control group. Mitotracker DR and live/dead aqua staining was performed at room temperature and the samples run on a Dako Cyan flow cytometer with analysis by FlowJo software. Live cells were gated on those that had divided exactly once by CTV dilution. **G**: A similar experiment to that in B-D was set up except that a DC + peptide stimulated group was treated with rapamycin, and the ImageStream analysis of uropodium area was performed as described in the methods section.

Purified CD4^+^ T cells were cell trace violet (CTV) labelled and stimulated in vitro with bone marrow derived dendritic cells (bmDC) and their cognate antigen (Dby) peptide (21, 31). We compared their responses to an antigen presenting cell free stimulation by anti-CD3 plus anti-CD28 coated beads. At the end of the primary culture, cells were stained for mitochondria (Mitotracker DR), with conventional antibody labelling for cell surface CD4, plus intracellular staining for pS6 as an indication of mTOR activation. We used both conventional flow cytometry and imaging flow cytometry for analysis, with similar results, to determine the optimal conditions for both mTOR activation and to identify conditions where we could observe a similar bimodal distribution of CD4, mTOR activation and mitochondria to that previously described (9, 13, 38). Although standard RPMI+10% FCS culture medium initially supported these observations, we found poor reproducibility especially with different serum batches. For this reason, we moved to a more defined medium formulation (Advanced RPMI) before further optimisation. An example optimisation experiment is shown in Figure 2B-D which examines some of the requirements of the tissue culture conditions, in this case titrating the concentration of FCS, together with the addition of both IL-2 and TGFβ Section 3.10 gives further details on the role of these two cytokines. Analysis was performed on day 3 (before nutrient depletion and intrinsic mTOR inhibition take place). Under these conditions, 1% FCS (as indeed recommended by the medium manufacturer) was optimal for both bimodal mitochondria staining (Figure 3C) and maximal pS6/mTOR activation (Figure 2D). This required bmDC/Dby peptide stimulation (at a previously determined optimal concentration of 100nM for A1RAG T cells) while such bimodality was never observed with anti-CD3/CD28 beads under any conditions tested. Note that while TGFβ suppressed proliferation after CD3/CD28 stimulation (39), its addition gave a more reproducible bimodal mitochondria staining with improved cell proliferation and survival when T cells were stimulated with bmDC plus antigen. The addition of TGFβ also allowed us to compare fate decisions of conventional and regulatory T cells subsets.

**Figure 3:**
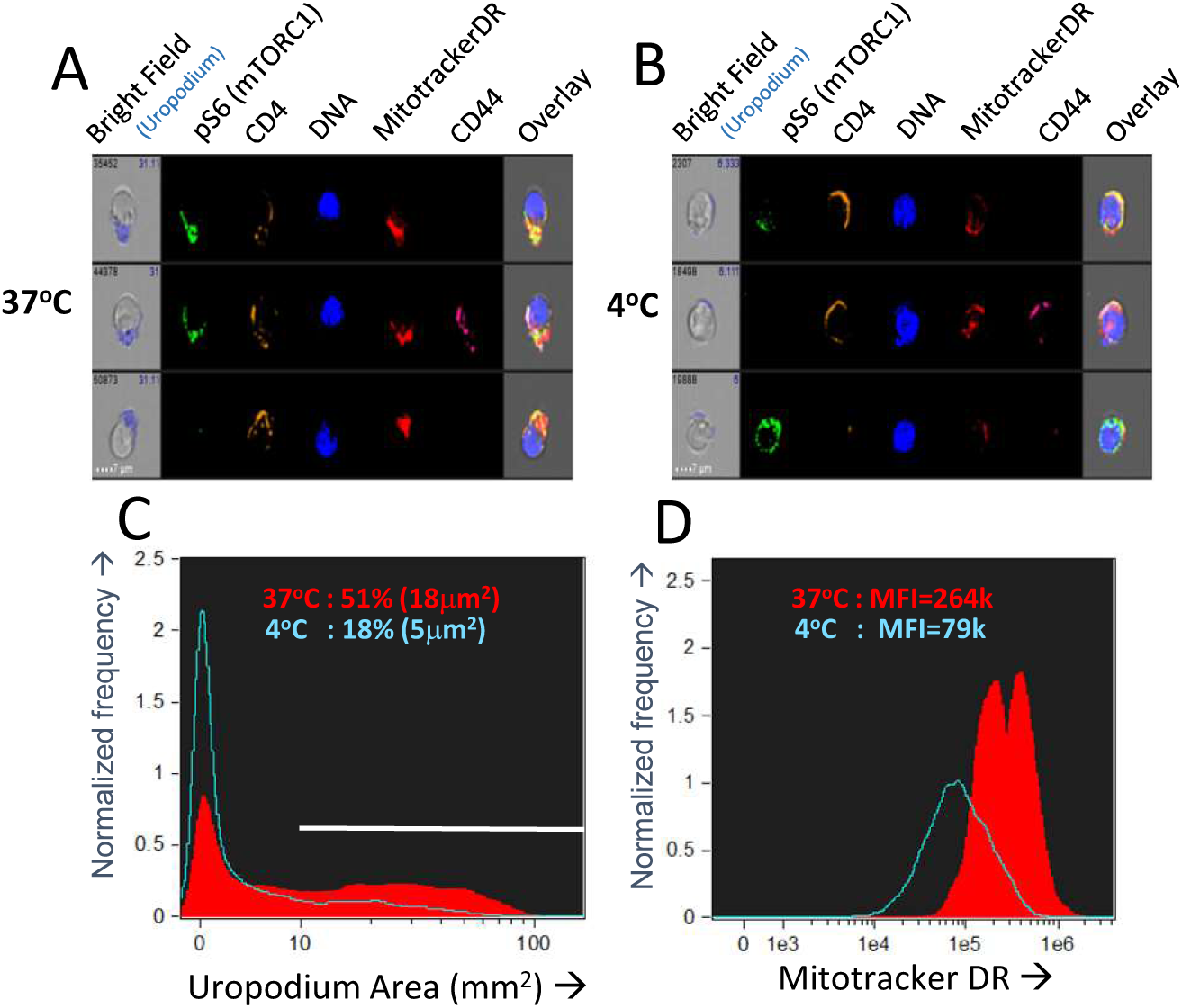
Optimising fixation and staining methodology. **A-D**: CTV-labelled naïve female A1RAG CD4^+^ T cells were stimulated with bmDC + 100nM Dby peptide with IL2, TGFβ and ATRA. Staining and fixation was performed in two different ways: either the “in situ” method at 37°C was used (**A** and red filled lines in **C, D**) or cells were conventionally harvested, spun down and labelled in PBS+1%BSA with fixing and permeabilisation all at 4°C (**B** and blue lines in **C, D**). Cells were gated and uropodia area determined as described in the methods. One of two similar experiments shown.

The bimodal mitochondrial distribution was found to be dependent on robust mTOR activation, as rapamycin (mTORC1 inhibitor), Torin 1 (an inhibitor of total mTOR ATP-dependent activity) and amino acid starvation (30) all reduced the bimodality towards a uniform single peak (Figure 2E-F). The proportion of cells with uropodia and the uropodium area were also reduced by mTOR inhibition, although not completely (Figure 2G).

### 3.2 Optimisation of fixation and staining for imaging flow cytometry

Even with this optimised in vitro culture system, we still encountered issues of reproducibility in the staining. We were unable to find any cells with the very distinctive morphology of telophase – although we found images with some features of late cytokinesis, but these could not be reliably distinguished from doublets and conjugates. Similarly, although we often identified cells with uropodia, the reproducibility between experiments was poor. We reasoned that both of these structural features were dependent on dynamic metabolic processes that were being disrupted during cell harvesting and staining. For this reason, we developed an “in situ” staining and fixation method (see methods). Figure 3 A-D compares this “in situ” staining/fixation at 37°C with conventional harvesting and cell staining (Mitotracker DR at room temperature then cell surface staining at 4°C), showing that only the “in situ” method reliably maintains both uropodia and clear bimodal mitochondrial staining. Similarly, late anaphase and telophase cells were readily identifiable if the cells were fixed/stained at 37°C in situ (see methods and Figure 1), but not with conventional staining at 4°C (not shown).

### 3.3 A CD4^+^ T cell memory response in vivo is associated with a bimodal distribution of mitochondria

Our in vitro optimisation focussed on the bimodal mitochondrial distribution in A1RAG CD4^+^ T cells responding to antigen, but we needed to check if this remained relevant to in vivo responses. We analysed draining lymph nodes from female A1RAG mice in mice undergoing a secondary rejection of male skin grafts, where we reasoned we should see high frequencies of both effector and memory CD4^+^ T cells. We compared this to the lack of rejection in mice previously rendered tolerant of male skin (31). Figure 4A shows the experimental design for these transplantation experiments. We had previously found very few differences in the proliferative responses or cell surface phenotypes of CD4^+^ T cells from the draining lymph nodes of both rejecting and tolerant mice (22, 40), and here we also observed similar levels of CD4 T cell activation, as indicated by high CD44 expression (Figure 4B, C). There were, however, striking differences in the cellular distribution of mitochondria (Figure 4D). While T cells from tolerant mice had uniform numbers of mitochondria per cell, rejecting T cells had a highly reproducible bimodal distribution with two clear populations differing by approx. 5-fold in their Mitotracker DR staining (Figure 4D). While this confirmed the in vivo relevance of the bimodal mitochondrial staining, the graft microenvironment and its interaction with the immune response (31, 40) is very difficult to manipulate experimentally, and harvesting, purifying and staining cells risks destroying important information related to their function, such as their nutrient status, signalling, and structural properties (as shown above). We therefore returned to the optimised in vitro culture system for further mechanistic investigations.

**Figure 4:**
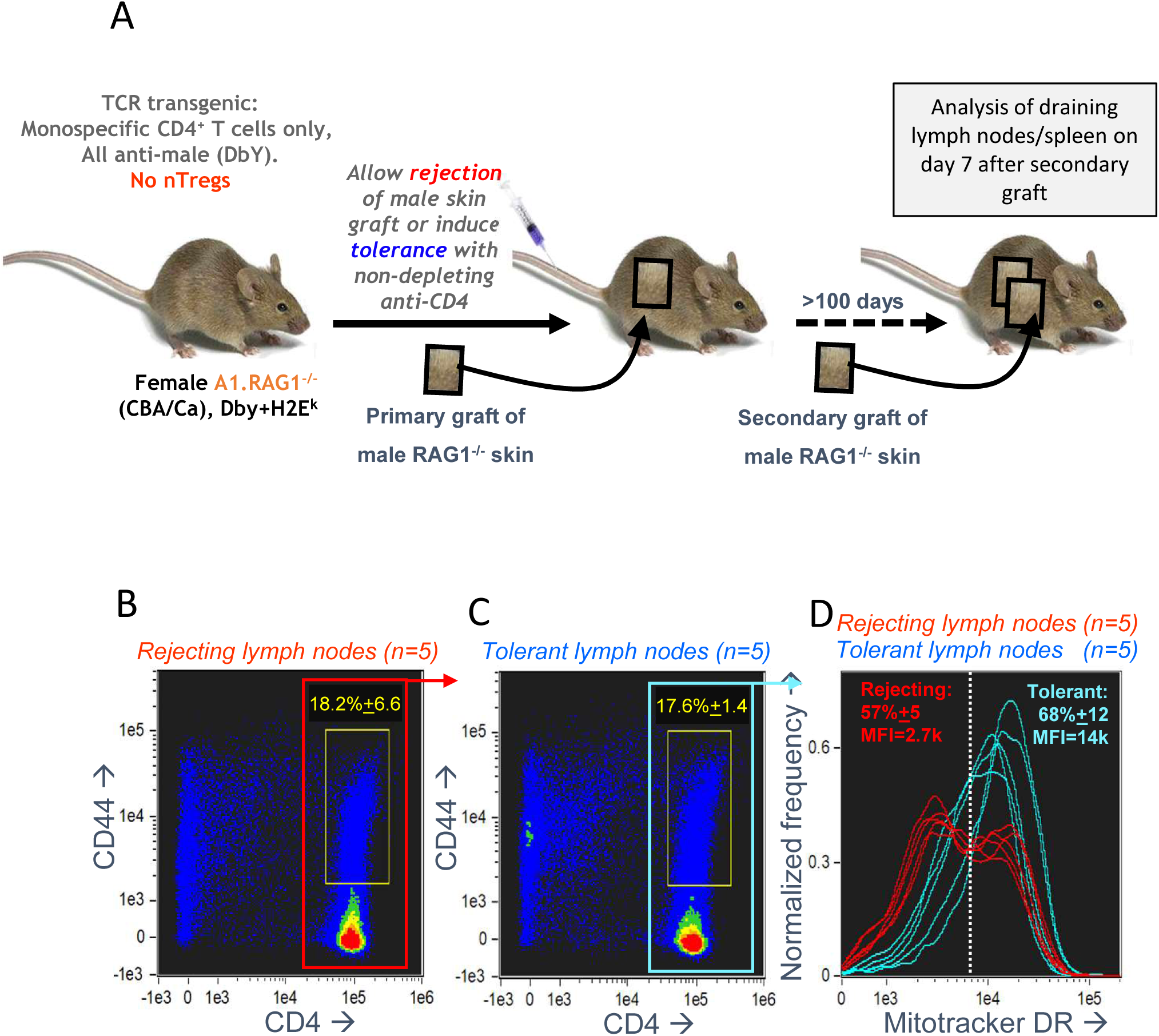
Antigen specific CD4^+^ T cells undergoing a secondary graft rejection response have a bimodal distribution of mitochondria. **A**: Schematic of the in vivo model used to compare secondary memory responses of antigen specific CD4^+^ T cells to a challenge skin graft after rejection or the induction of tolerance. **B-D**: Draining lymph nodes were taken, 7 days after a secondary challenge with male skin, from female A1RAG mice that had been either been allowed to reject a male skin graft or had been made tolerant by anti-CD4 treatment (5 mice per group). Cells from each mouse were individually stained with Mitotracker DR, CD4-PE-CF594, CD44-APC-eflour780, and 7AAD and 20,000 images were acquired by ImageStream for each sample. Singlet, viable cell images were gated on 7AAD intensity (2N DNA) and high aspect ratio, bright field area and low contrast. All rejecting (**A**) and tolerant (**B**) gated images are pooled and shown in the plots, and the percentage (mean±SD) of activated CD4^+^CD44^+^ T cells (within yellow gates) are indicated. The Mitotracker DR intensity of individual rejecting and tolerant CD4^+^ gated cells (red [**B**] and blue [**C**] boxes) is plotted in **D**.

### 3.4 Two cell fates distinguished by their differential expression of mitochondria, CD4 and uropodia

We first tested whether the two populations with high or low mitochondrial staining were transient and interconvertible, or represented two different cell fates. We did this by tracking the inheritance of mitochondria pre-labelled with Mitotracker DR (which covalently labels proteins within active mitochondria), that is then diluted as mitochondria partition into the daughter cells after division, as shown in Figure 5A, B. By additionally labelling for total mitochondria with a different dye (Mito-ID-red) after fixation at the end of the experiment, we could determine whether the cells with high or low mitochondria were generated by a differential inheritance (ie. whether there was any deviation between Mitotracker DR and CTV dilution during cell proliferation), and whether there was any mixing or interconversion between the two populations. The high Mito-ID-red staining (total mitochondria) population, which were also those with large uropodia (Figure 5E), showed a regular two-fold dilution of the Mitotracker DR stain with each cell division (Figure 5F). In contrast, the population with low Mito-ID-red staining did not develop significant uropodia, showed a considerable loss of Mitotracker DR staining even before the first cell division, and thereafter continued regular two-fold dilutions from this low level as cells proliferated (Figure 5G). This demonstrates that two independent cell fates had been generated even before the first cell division that can be distinguished readily by the numbers of mitochondria they possess, and this difference was maintained and inherited through subsequent cell generations.

**Figure 5:**
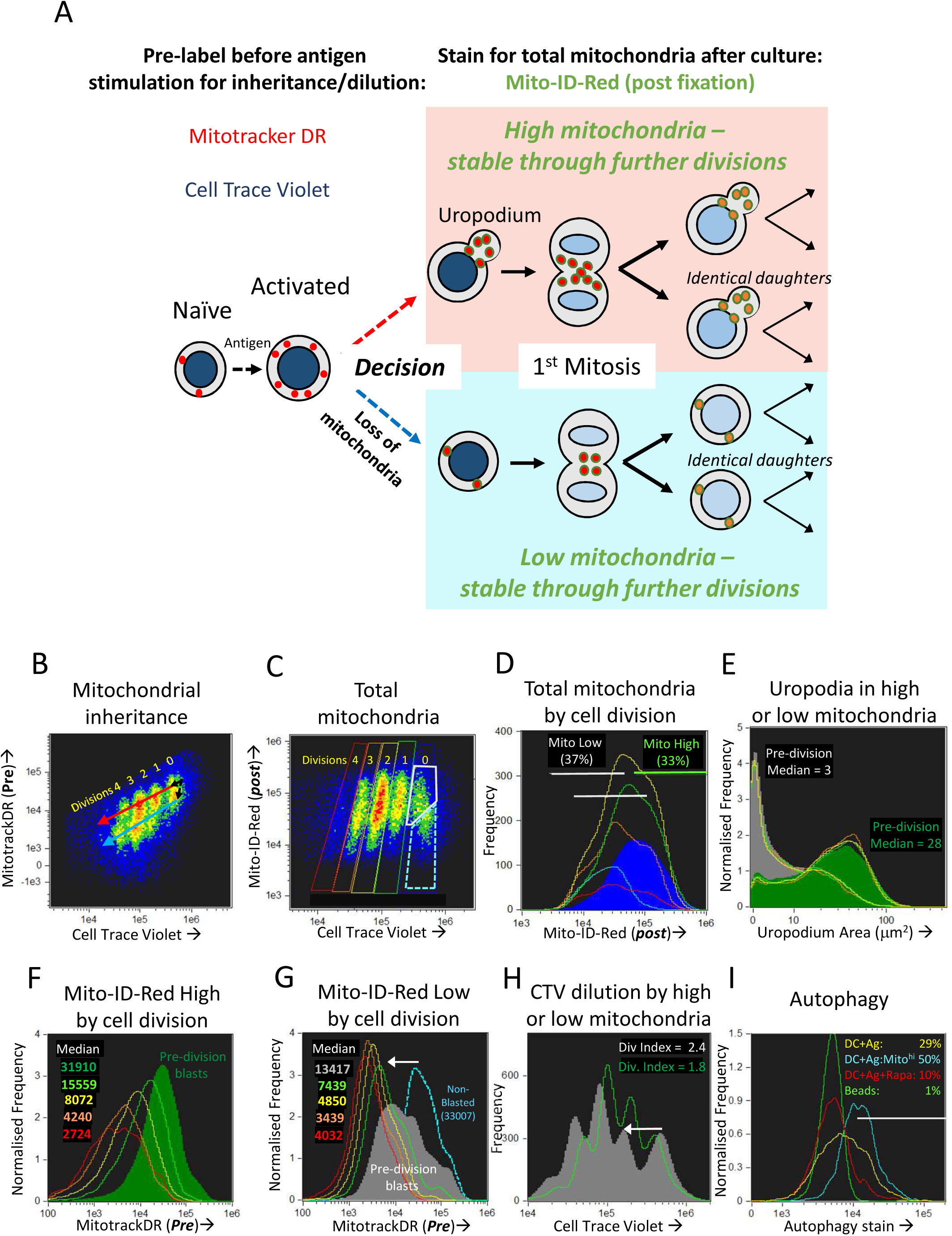
The bimodal distribution of mitochondria represents two lineages of proliferating cells that either do, or do not, develop uropodia before the first cell division. A: Explanatory diagram showing how pre-labelling with both Cell Trace Violet and Mitotracker DR allows the tracking of mitochondrial inheritance with cell division, while post-labelling with Mito-ID-Red indicates the total number of mitochondria per cell at the end of the culture, which shows that two separate lineages of cells proliferate in parallel and do not interconvert. **B-I**: Naïve female A1RAG CD4^+^ T cells were labelled with both CTV and Mitotracker DR and stimulated with bmDC + 100nM Dby peptide with IL-2, TGFβ and ATRA, for 3 days. Cells were labelled “in situ” with CD4-APC-Cy7 and live/dead aqua, fixed and permeabilised, then stained with Mito-ID-Red and Sytox green, and 30,000 images acquired by ImageStream. All images were gated for live, singlet, G_0_/G_1_ and CTV^+^ cells. Panel **B** shows the dilution of Mitotracker DR with each cell division by CTV dilution. Panels **D-G** show histograms of the indicated parameters gated in C for each cell division (0=Blue, 1=green, 2=yellow, 3=orange, 4=red) with pre-division activated blast cells (high mitochondria [grey box] and bright field area >90μm^2^) and non-blasted cells (low mitochondria [dashed blue box] and bright field area <90μm^2^). Each population is further gated in panel **D** for high (green) or low (grey) total mitochondria (Mito-ID-Red stain). Panel **E** shows that the Mito-ID-Red high and low populations have either large (>10μm2) or small/no (<10μm^2^) uropodia respectively, with very similar proportions of each across all cell divisions. While the Mito-ID-Red high populations show a regular 2-fold dilution of the pre-stained Mitotracker DR (**F**), the Mito-ID-Red low gated cells lose most of their Mitotracker DR staining during their activation from non-blasted to pre-division blast cells (panel **G**, white arrow), with regular 2-fold dilutions to a background of 4000 after that. CTV was also lost at the same time point (panel **H**, white arrow), but returned to regular 2 fold dilutions thereafter. One of two similar experiments shown. In a separate experiment (**I**), CTV labelled A1RAG CD4^+^ T cells were stimulated, in the presence of IL2, TGFβ and ATRA, with bmDC + Dby peptide, with (red line) or without (yellow line) rapamycin, or with CD3/CD28 beads (green line). Cultures were labelled at different time points (only 48h shown) in situ, with autophagy green detection reagent (Abcam: 1/2000), Mitotracker DR, CD4-PECF594 and CD44-APC-Cy7 for 40 mins, then fixed with 2% formaldehyde at 37°C for 15 mins. After a single wash in PBS + 1% BSA, samples were immediately run on the ImageStream. Images were gated on focussed, live, single CD4^+^ cells with undiluted CTV staining. One of 3 similar experiments is shown.

Those cells without uropodia, which had also lost mitochondria before the first cell division, also lost some of their CTV staining at this same point (Figure 5H). One explanation for this combined non-specific loss of two different covalent protein stains in cells destined for just one of the two cell fates, and then only before the first cell division, might be a brief period of autophagy/mitophagy (41). Inhibitors of autophagy (chloroquin or spautin 1) compromised all T cell activation and proliferation, consequently blocking any opportunity to observe the two cell fates (not shown), but we were able to detect increased staining with an autophagy dye from 48h after stimulation, and before cell division, in the DC + Ag group (Figure 5I). This represented about half of the cells that had increased both their size (ie. blasted) and mitochondrial mass since first stimulated, but those cells destined to become memory cells showed higher levels of autophagy associated with a subsequent loss of mitochondria (and Mitotracker DR) as well as other cellular proteins (CTV label) before the first cell division.

### 3.5 No role for asymmetric cell divisions in generating two cell fates

When naïve T cells make an apparent binary cell fate choice during an asymmetric cell division (Figure 6A), the two daughter cells are characteristically CD4 (or CD8) high versus low (9), and differ in their numbers of mitochondria (38). The two populations generated correspond to short-lived effector versus long-term memory cells (9, 11). Controversially (6, 37, 42), it is claimed that the binary nature of this fate decision is the result of an asymmetric inheritance of various transcription factors (8, 10) and signalling components that drive these diverging cell fates (13, 14). Effector and memory T cells also differ in their PI3k/mTOR signalling and metabolic profiles (12, 13, 43), although uropodium development has not previously been reported in this context. Our initial experiments had also appeared to support this asymmetric model, but during the optimisation of staining and fixation methods we realised that we had likely been misled by technical artefacts. After optimisation and flow cell imaging analysis, we could objectively and unambiguously identify (see Figure 1H-N gives the masking and gating strategy) all the images of rare cells in late anaphase and telophase (without the use of any mitotic inhibitors) and accurately measure whether any cell markers were polarised towards one daughter cell or the other (Figure 6C-O). Multiple experiments confirmed that all uropodia were lost after prophase and that there was no significant polarisation of any of the cell surface markers (CD4, CD44, CD25: Figure 6C, D, E), transcription factors (Tbet, RORγt, Foxp3, IRF4, NFκB: Figure 6F-J) nor mitochondria (Figure 6N) at telophase, in neither the first, nor subsequent cell divisions, effectively ruling out any role for asymmetric cell divisions in our system. This is consistent with the data of Figure 5 and suggests a model where the decision to develop an effector cell fate (with uropodia development) takes place before the cells enter their first division, and the two cell lineages then continue to proliferate in parallel (Figure 6B).

**Figure 6:**
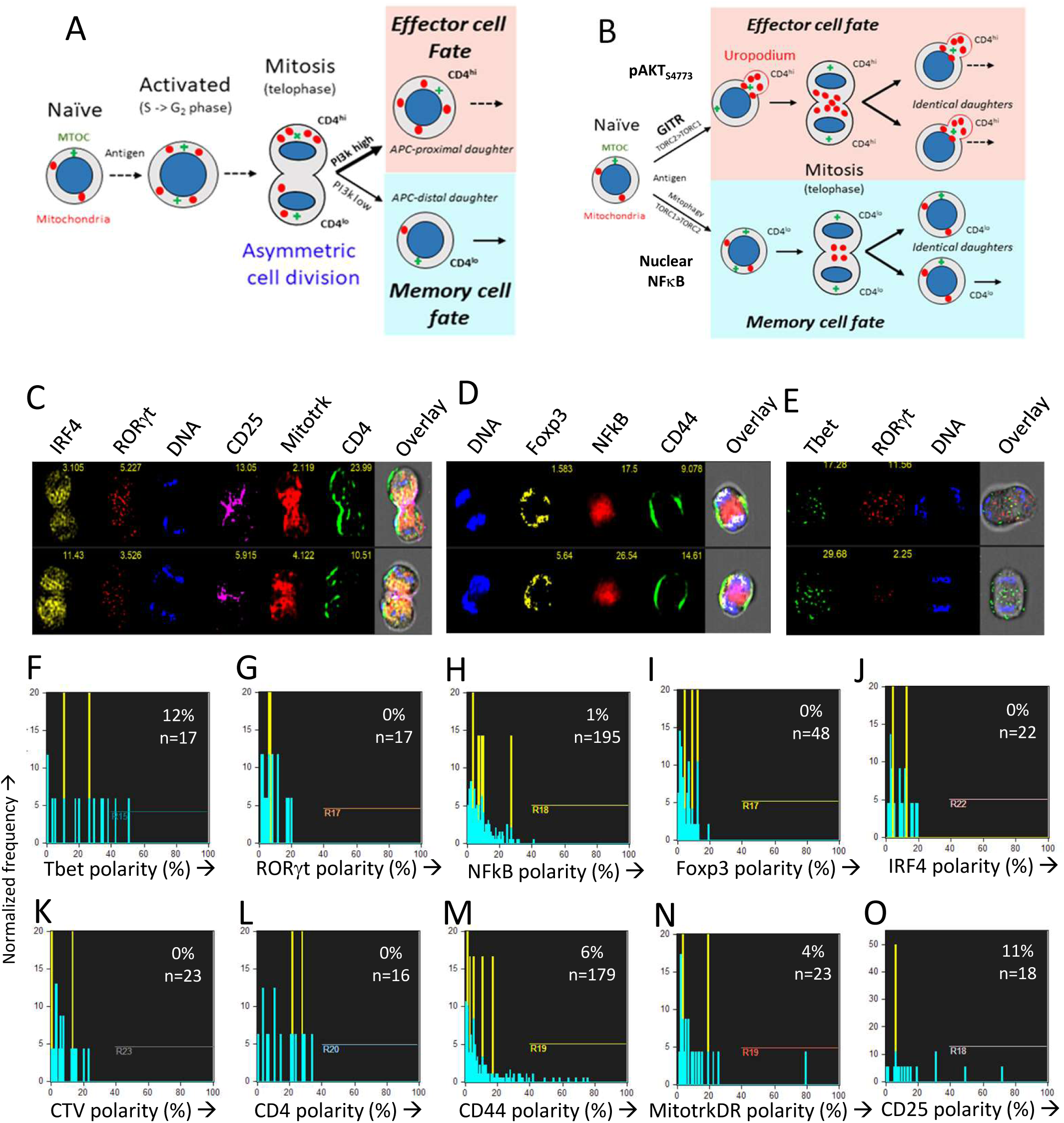
The CD4^+^ T cell fate choice associated with uropodium development does not depend on asymmetric cell divisions. **A**: Depiction of a binary cell fate decision as a result of an asymmetric first cell division. The effector and memory cell fates result from a differential inheritance of mitochondria, CD4 and PI3k signalling between the two daughters. **B**: A stochastic cell fate decision to develop uropodia during initial activation and before any cell division. The chance of any individual cell becoming either an effector cell, and developing uropodia, or a memory cell without uropodia, depends on the balance of a number of interacting signalling pathways (eg. GITR, mTORC1, mTORC2, NFκB) during its initial activation. Symmetrical cell divisions can then take place regardless of the cell fate decision taken. **C-O**: CTV labelled naïve CD4^+^ T cells from female A1RAG were stimulated with bmDC + 100nM Dby peptide with IL-2, TGFβ and ATRA for 48 h (when the average number of cell divisions was only 0.7). No mitotic inhibitors were used. Cultures were labelled “in situ” with (**C, L**) CD4-APC-Cy7 and CD25-BV605 or (**D, M**) CD44-APC-eflour780, plus live/dead aqua and Mitotracker DR (**C, E, N**), and fixed at 37°C by addition of 40% formaldehyde to 10% v/v for 15 minutes. The medium and fixative were then carefully aspirated and fix/perm buffer added for 2 hours at 37°C. Intracellular stains used were (**C** and where indicated): IRF4-PE, RORγt-PE-CF594, 7AAD or (**D** and where indicated): Sytox Green, Foxp3-PE-Cy7, NFκB-p65-APC or (E and where indicated): Tbet-Alexa488, RORγt-PE-CF594, 7AAD followed by ImageStream analysis. Telophase and late anaphase cells were automatically gated and all identified cells expressing the marker of interest are included in the histograms of polarity scores (with numbers indicated). Cells in their first mitosis (ie. undiluted CTV) are indicated in yellow. Data shown is from ~600,000 images obtained by pooling 3 independent experiments (under standard, Th1 or Th2 cytokine conditions) and is representative of at least 3 similar datasets.

### 3.6 Confirmation that the cells with high mitochondria have uropodia

Up to this point, our identification of uropodia depended on an image mask which measures the area of any single, large protrusion beyond the regular, circular shape of the cells (2D image). It is known that functional uropodia are required for normal immune responses (28,44) and that this is associated with a specific organisation of the cell surface and cytoplasmic organelles, as summarised in Figure 7. Uropodia are also important for cell migration, when they can be found at the rear of motile lymphocytes. Mitochondria localise to the uropodium by moving along microtubules that originate from the microtubule organising centre (MTOC) at the base of the uropodium. Mitotracker DR staining was indeed localised to the rear facing uropodia by live cell video imaging of CD4^+^ T cells being activated by DC + Dby peptide (Supplementary Video 1).

**Figure 7:**
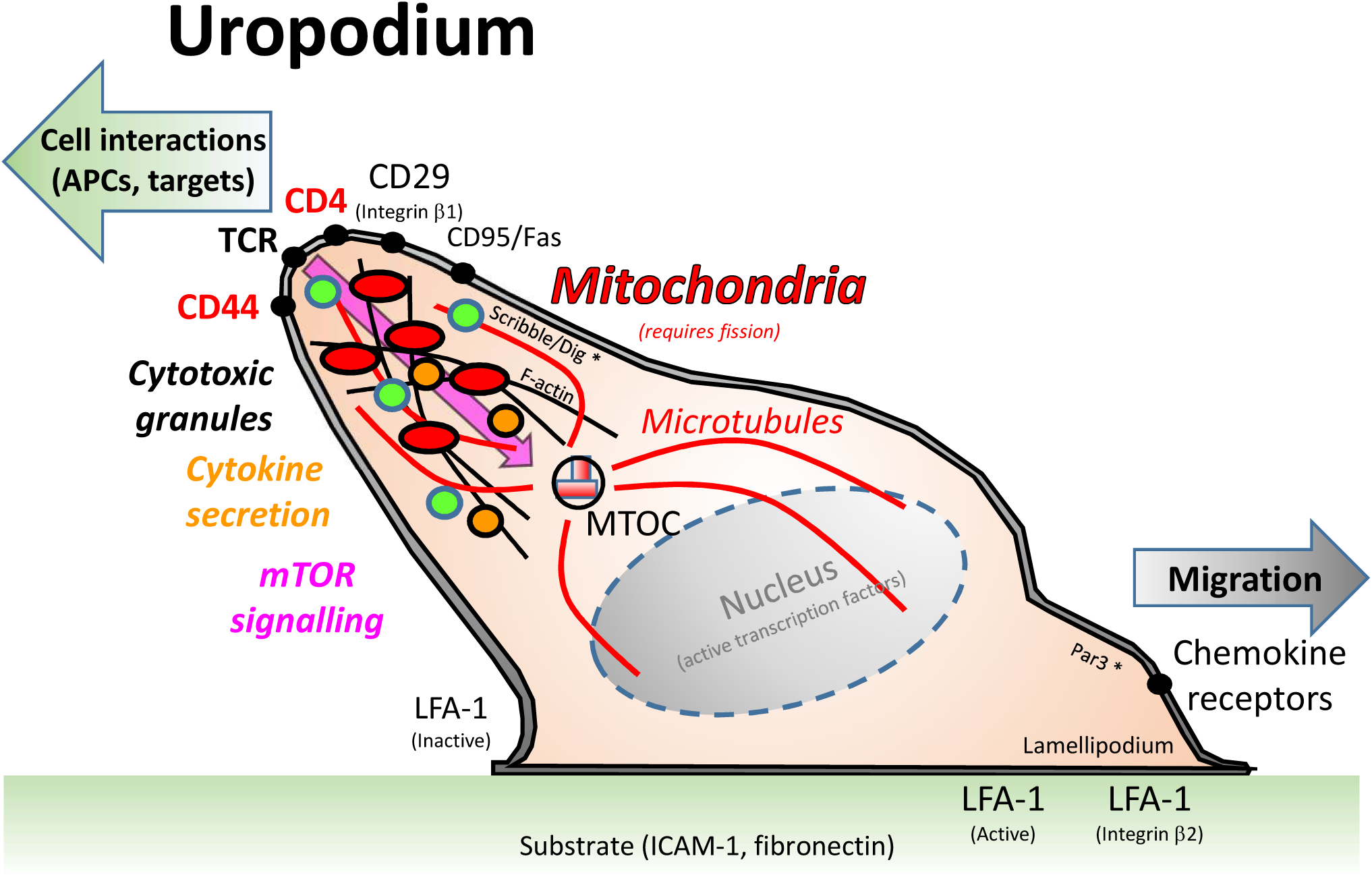
A depiction of the uropodium structure in lymphocytes. Adapted from (44). Uropodia are found at the back of activated lymphocytes that are migrating on a substrate, usually an intracellular matrix containing integrin ligands such as ICAM-1 or fibronectin, towards a chemokine cue. The uropodium is organised as a large, finger-like projection by the cytoskeleton and microtubules, with the microtubule organising centre (MTOC) at its base. The polarisation of the cell is maintained by the Scribble/Dig and Par3 complex, which are the same components that have been claimed to be needed for asymmetric cell divisions (10). It is thought that the uropodium is responsible for the interaction of T cells with other cells, such as antigen presenting cells and targets of cytotoxicity, and therefore expresses high levels of TCR, CD4/8 and relevant adhesion molecules (CD44, CD29). The cytotoxic granules of effector T cells are also contained within the uropodium, and cytokines are secreted from them. Mitochondria, and many other organelles are also concentrated within the uropodium, which contains the bulk of the cell cytoplasm.

**Supplementary Video 1:**
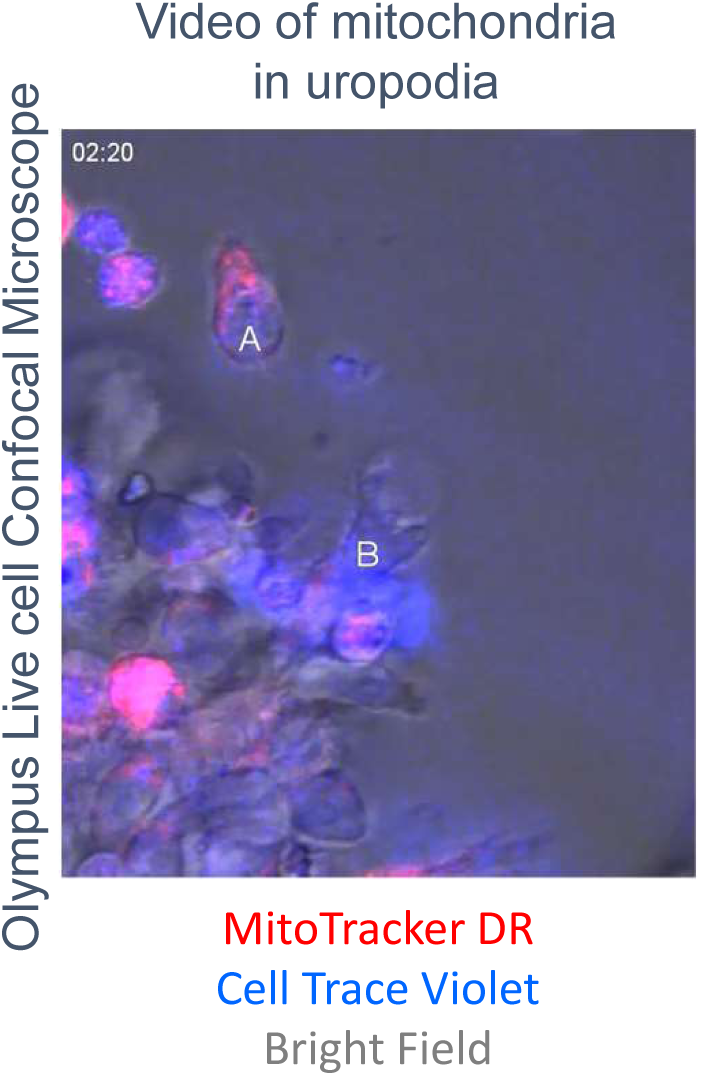
Live cell imaging of migrating, cell trace violet labelled CD4^+^ T cells (A, B) with mitochondria (Mitotracker DR stained) within uropodia at the rear.

We confirmed that the structures we identified on the high mitochondrial staining cell population were indeed uropodia by the statistical analysis of large numbers (10-50,000 per sample) of antigen-stimulated CD4^+^ T cell images for their shape and the expression and localisation of other cell components (Figure 8A-N). More than half of all the uropodia showed clear staining at their base for the MTOC (γ-tubulin: Figure 8A, C), and were high in CD44, 37% of which, on average, was located on the uropodium surface (Figure 8F, K). Uropodia also contained 50% of all the mitochondrial staining (Figure 8E, J). In the cells with uropodia, CD4 expression was also much higher and localised to the uropodium surface (Figure 8G, L). As shown above, uropodium development was associated with robust mTOR activation and both mTORC1 (pS6) and mTORC2 (pAKT_S473_) (45, 46) levels were higher in the cells that developed them (Figure 8H, I), with the latter mostly localised within the uropodium (Figure 8N).

**Figure 8:**
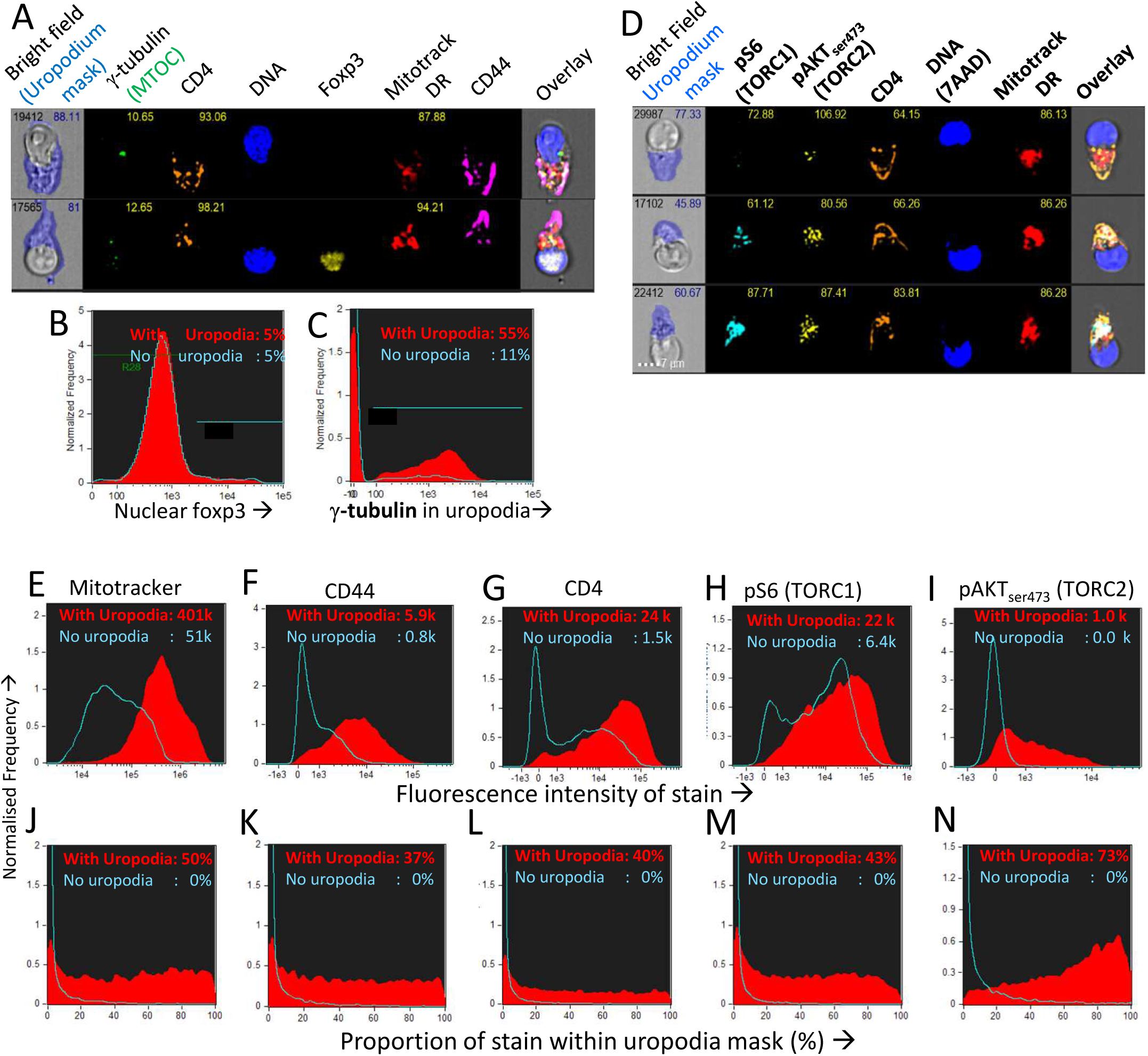
Confirmation that the image masking strategy used is correctly identifying uropodia containing high numbers of mitochondria, high CD4 and CD44 expression, and mTOR signalling. **A-C**: CTV-labelled naïve female A1RAG CD4^+^ T cells were stimulated with bmDC + 100nM Dby peptide with IL2, TGFβ and ATRA for 3 days. Cells were labelled “in situ” with CD4-PE-CF594, CD44-APC-Cy7, Mitotracker DR, and live/dead aqua, fixed and permeabilised, and intracellularly stained for γ-tubulinAlexaFluor488, and foxp3-PE-Cy7. Example images are shown in **A**. The distribution of nuclear foxp3 staining was identical comparing cells with or without uropodia (**B**) and γ-tubulin staining, indicating the MTOC was strongly associated with uropodia (**C**). **D-N**: CTV labelled naïve A1RAG CD4^+^ T cells were stimulated with bmDC + 100nM peptide under optimised conditions (see methods) for 3 days and then “in situ” labelled with CD4-PE-CF594, CD44-APC-eflour780, CD62L-BV605 (not shown), live/dead aqua and Mitotracker DR, fixed and permeabilised, then stained with pS6-Alexa488, pAKTS473-PE, Foxp3-PE-Cy7 (not shown) and 7AAD for ImageStream analysis. All images were gated for live, singlet cells in G_0_/G_1_ as previously. Example images, with uropodia masks indicated, are shown in **L**, and cells with a uropodium area greater or less than 10μm^2^ were defined as positive (red filled histograms) or negative (blue histograms), respectively. **E-I** show the intensity histograms for each stain of interest, while **J-N** show the proportions of each stain that fall within the uropodium gate for each image. Median values for each plot are indicated. Shown are representative examples of 3 or more independent experiments.

### 3.7 Cell fate choice is stochastic and dependent on both mTORC1 and mTORC2 signalling

We found that across a wide range of Dby peptide concentrations (Figure 9A, B; only two extremes shown), two distinct populations of cells developed, one bearing uropodia and the other lacking them, based on their measured area (Figure 9F, G). The fact that we observed two discrete populations of cells (either with or without uropodia rather than a continuum of increasing uropodium size) suggests that individual T cells were still making a binary fate decision, but in a manner that was stochastic and not determined by cell division. The chance of an individual T cell developing an uropodium seemed to depend on the strength of signalling through the mTOR pathway (as shown above, Figure 2G) which led us to seek evidence of discrete signalling states within this pathway that were associated with uropodium development. Mathematical models suggest mechanisms by which such discrete and stochastic signalling states may arise without pre-existing heterogeneity (47). When we simultaneously stained for both mTORC1 (pS6) and mTORC2 (pAKT_S473_) signalling (48), we reproducibly found a total of 6 distinct populations: 3 with weak/negative, intermediate or high mTORC1 staining, differing from each other by an order of magnitude, with each of these 3 populations further split into either mTORC2 positive or negative cells (Figure 9A-E). Cells with uropodia were found predominantly within the mTORC2 positive population that were mTORC1 intermediate (Figure 9F, G). The distribution of all CD4^+^ T cells across these 6 populations depended on the concentration of antigen/TCR stimulation, which mainly increased mTORC1, while mTORC2 signalling required antigen presenting cells (bmDC: Figure 9C, D). Further analysis of the localisation of pAKT_S473_ confirmed that it was specifically mTORC2 signalling that was localised to the uropodia rather than total AKT (Figure 9K, L, M) or PI3K signalling through pAKT_T308_ (not shown). The bmDC could provide some mTORC2 signalling independent of TCR stimulation (Figure 9E). We also found that total NFκB p65 was strongly up-regulated with DC + antigen (Figure 9Q) in all cells, but a specific increase in the nuclear localisation of NFκB, indicating signalling, was highest in the cells lacking uropodia (Figure 9R). Note that NFκB signalling is thought to be important for memory cell differentiation and maintenance (49). CD3/CD28 bead stimulation, by comparison, was poor at upregulating total or nuclear NFκB (Figure 9Q, R), despite inducing strong cell proliferation (Figure 9P).

**Figure 9:**
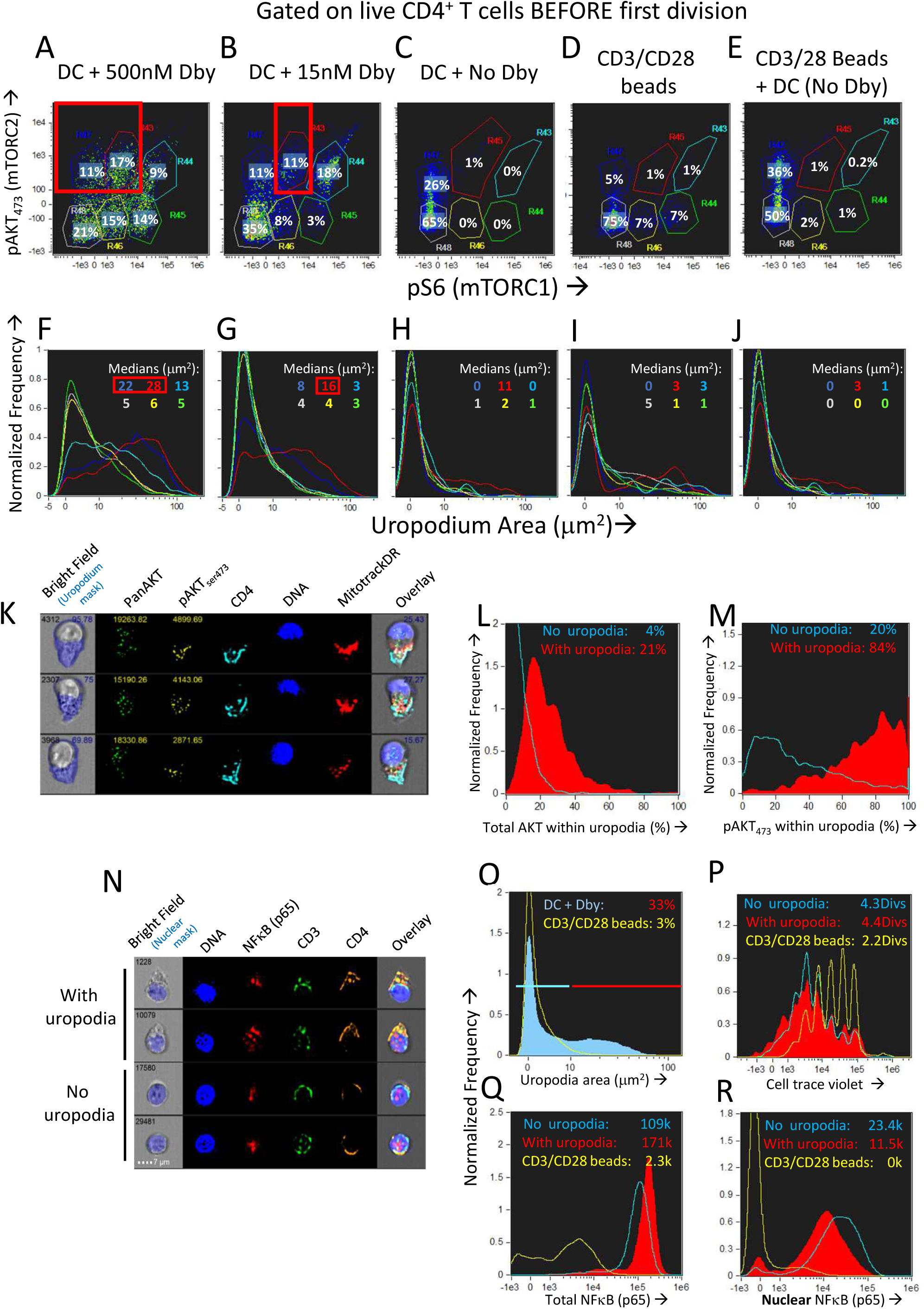
Development of uropodia is associated with strong pAKT473/TORC2 signalling, intermediate pS6/TORC1 activity and low NFκB signalling. **A-J**: CTV labelled female A1RAG CD4^+^ T cells were stimulated for 3 days in IL-2, TGFβ and ATRA, with bmDC while the Dby peptide was titrated from 500nM down to zero (examples shown in **A-C** and **F-H**) or CD3/CD28 beads were used as stimulation either alone (**D, I**) or together with DC but no Dby peptide (**E, J**). Cultures were labelled “in situ” with CD4-PE-CF594, CD44-APC-eflour780, Mitotracker DR and live/dead aqua, fixed and permeabilised, followed by staining with pS6-Alexa488, pAKT_S473_-PE and 7AAD for ImageStream analysis. Gating was for live, singlet cells in G_0_/G_1_ that had not diluted their CTV (ie. before any cell division). The uropodia area distributions of the six populations gated in panels **A-E** are colour-coded and shown in the histograms of panels **F-J** respectively. Large uropodia were only induced in cells stimulated with bmDC plus Dby peptide, and where pAKT_S473_ was high and pS6 was simultaneously intermediate or low (red boxes). One of 3 similar experiments shown. **K-M** shows a similar experimental set up (1 of 2) except that a pan-AKT-Alexa488 antibody was used in combination with the pAKT_S473_-PE staining, showing that total AKT was not restricted to the uropodia (example images in **K** and histogram in **L**) while pAKT_S473_, indicating signalling, was uropodia restricted (**M**). Median values for % within uropodia are shown. **N-R**: The experiment shown (1 of 3) used either bmDC + 100nM Dby peptide or CD3/CD28 bead stimulation, was labelled “in situ” with live/dead aqua and CD25-BV605 (not shown), fixed and permeabilised, then stained intracellularly with CD4-PE-CF594, CD3-APC-Cy7, NFκB-p65-APC, (Foxp3-PE-Cy7, CD95-FITC, not shown) for ImageStream analysis. Example images of cells with and without uropodia (stimulated by bmDC + Dby) are shown (**N**) while all live, singlet G_0_/G_1_, DC+Dby (filled blue) or CD3/CD28 bead (yellow) stimulated cells are shown in the histogram of uropodium area (**O**). Histograms of the CTV dilution profiles of bmDC stimulated cells either with (filled red histograms) or without (blue lines) uropodia, or CD3/CD28 bead stimulated cells (yellow lines) are shown in **P** with the mean number of cell divisions indicated. The intensity histograms for total NFκB-p65 (**Q**) or NFκB restricted to the nucleus (**R**) are shown with median intensity values indicated.

### Development of uropodia before the first cell division depends on mTORC2 signalling via GITR

We wondered which ligands on the DC might be providing the additional “co-stimulation” that could increase both mTORC2 and NFkB signalling compared to CD3/CD28 beads and at the same time, promote the cell fate decision and development of uropodia. It has been shown that some members of the TNFR family, ligands for which are known to be present on DCs (35), can signal via mTORC2 as well as through NFκB (46, 50). One member of the TNFR family that is upregulated rapidly upon T cell activation is GITR (TNFRSF18) while its ligand (GITRL; TNFSF18) is well expressed on bmDC (35, 51). GITR activation by its ligand, or by cross-linking with an agonistic antibody, acts to co-stimulate T cells at intermediate levels of TCR signalling (35). We found (Figure 10A-F) that agonistic GITR antibody coated on plastic together with intermediate concentrations of anti-CD3 (plus soluble anti-CD28) gave considerable enhancement of both nuclear NFκB (Figure 10E vs B) and mTORC2/pAKT_S473_ (Figure 10F vs C) together with increases in the number of cells with uropodia (Figure 10D vs A). Similar enhancements could be observed (Figure 10N-P) using antigen and bmDC stimulation and agonist GITR antibody in solution, where the rat IgG2b mAb (35, 36) can bind to Fc receptors on the APC for cross-linking. Furthermore, in this case where bmDC were already stimulating uropodium development, blocking of their GITRL gave a substantial loss of uropodia (Figure 10K vs H), mTORC2 signalling (Figure 10M vs J) and nuclear NFκB (Figure 10 L vs I) before the first cell division (yellow histograms), when the cell fate decision normally takes place (as shown above). With continued GITRL blocking, some uropodia did develop after the first cell division (and mTORC2 and NFκB signalling partially recovered) suggesting there may be redundancy for appropriate co-stimulation at later time points, for example with other members of the TNFRSF family known to alter the balance between effector and memory cells (5, 52).

**Figure 10:**
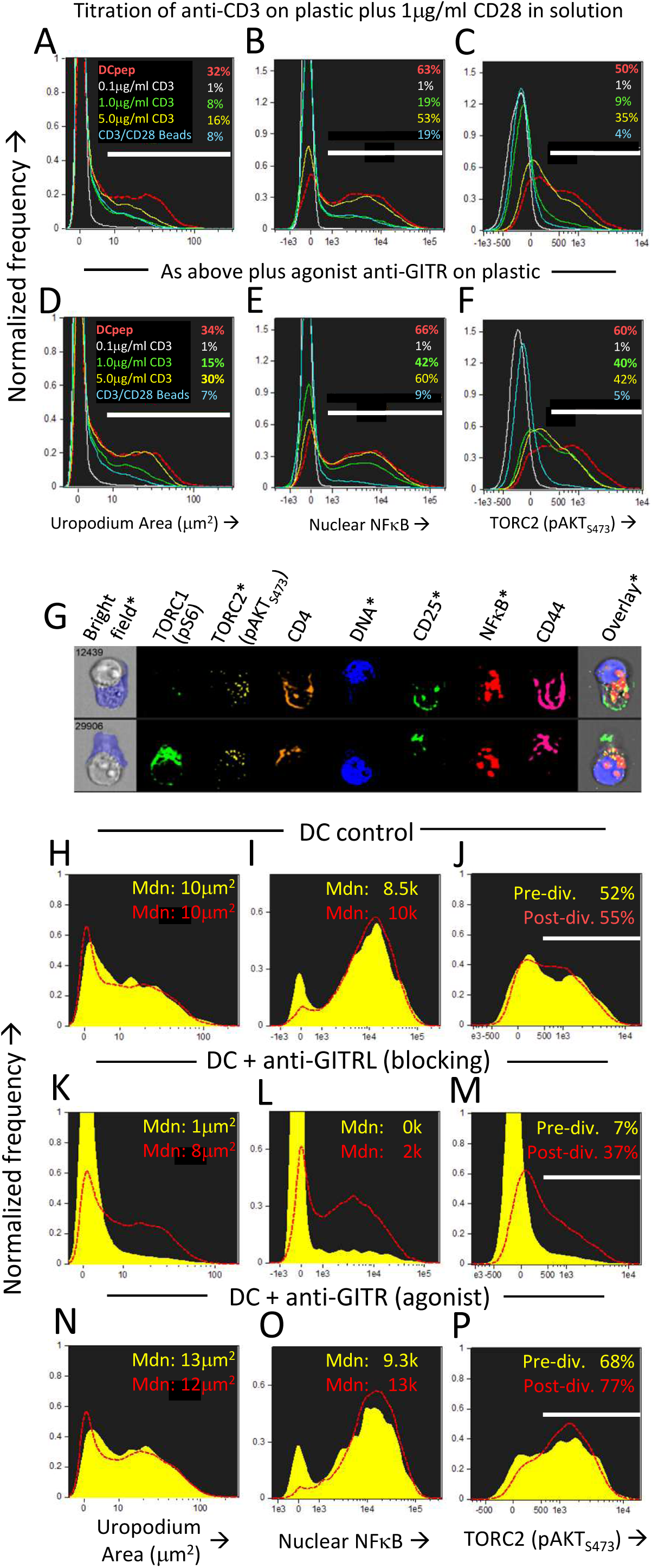
Uropodia development before first cell division depends on GITR signalling through mTORC2 and NFκB. **A-F**: CTV labelled female A1RAG CD4^+^ T cells were stimulated in the presence of IL-2, TGFβ and ATRA with either bmDC + 100nM Dby peptide (red, dashed histograms), CD3/CD28 beads (blue lines) or different concentrations (0.1μg/ml, white lines, 1.0μg/ml, green lines, or 5μg/ml, yellow lines) of anti-CD3 antibody coated on the tissue culture plastic, each concentration plus 1μg/ml anti-CD28 (37.51) in solution. In panels **D-F** an agonist antibody to GITR (YGITR 765.4) was also coated at 1μg/ml on the plastic. After 3 days, cultures were labelled “in situ” with CD4-PE-CF594, CD25-BV605, CD44-APC-eflour780 and live/dead aqua, fixed and permeabilised, followed by intracellular staining for pS6-Alexa488, pAKT_S473_-PE, NFκB-p65-APC, and 7AAD. Images for histograms shown were gated on live, singlet, CTV^+^, G_0_/G_1_ DNA content cells and the proportion (%) of cells that developed uropodia (>10μm^2^: **A, D**), stained for nuclear expression of NFκB (**B, E**) and pAKT_S473_ (**C, F**) are indicated. One of two similar experiments shown. **G-P**: An experiment similar to that above was set up, except that all cultures were stimulated by bmDC + 100nM Dby, either alone (example images in **G**, histograms in **H-J**), or with the addition of a blocking antibody to GITRL (YGL 386: **K-M**) or an FcR-binding, agonist antibody to GITR (YGITR 765: **N-P**), both at 10μg/ml in solution. Yellow filled histograms are gated on cells which have not divided (undiluted CTV) while dashed red histograms are gated on cells that have divided once or more. Median values are indicated. One of two similar experiments shown.

### 3.9 Uropodium development is associated with terminal effector and regulatory cell differentiation

We asked if one could map the development of uropodia (or not) to a similar effector versus memory T cell fate choice as previously claimed (9, 53) to result from asymmetric cell division? Effector T cells should proliferate rapidly, be able to migrate to the site of infection/inflammation, and express transcription factors, cytokines and cytotoxic molecules appropriate to their functional T cell subset (ie. Th1, Th2 etc) and, after terminal differentiation, die. These properties were indeed most clearly associated with the cells that developed uropodia (Figures 11 and 12), although migration is only implicit to uropodium function (44) as we did not test this directly. Naïve T cells stimulated to proliferate with CD3/CD28 beads did not develop uropodia, and showed almost no induction of effector T cell subset transcription factors by day 2 (Tbet, GATA3, RORγt, Foxp3: Figure 11A, D). Stimulation with antigen and bmDC induced these transcription factors, surprisingly in all possible random combinations and even before the first cell division, particularly in those cells that developed uropodia (Figure11B, C). These cells continued to proliferate (Figure 11D-F) and were functional as they also expressed similarly random combinations of cytokines (Figure 12A-F), suggesting that without any external selective pressure, this first wave of effector cells exhibit a diverse range of potential functions. A small proportion of the cells with uropodia were even co-expressing nuclear foxp3 together with effector cytokines (eg. Figure 12B), compatible with previous descriptions of foxp3^+^ regulatory cells with some Thelper subset properties (54). Granzyme B, important for effector cell cytotoxicity (55), was also expressed (Figure 12G, H), and tended to be localised to the uropodia (Figure 12I), as previously described (44, 56). Interestingly, where nuclear foxp3 was also present this localisation was reduced (Figure 12I). At later time points (day 7 of culture shown: Figure 12 J-M) there was evidence of apoptosis (increased bright field contrast) and necrosis (intracellular L/D Aqua staining, probably subsequent to apoptosis) in a population of cells which had achieved fewer divisions (average 4.7) than the viable cells (average 8.1: Figure 12 J). These expressed uropodia (Figure 12 K), more mitochondria (Figure 12 L) and higher levels of CD4 (Figure 12M), indicating that the effector cell lineage was short-lived under these conditions.

**Figure 11:**
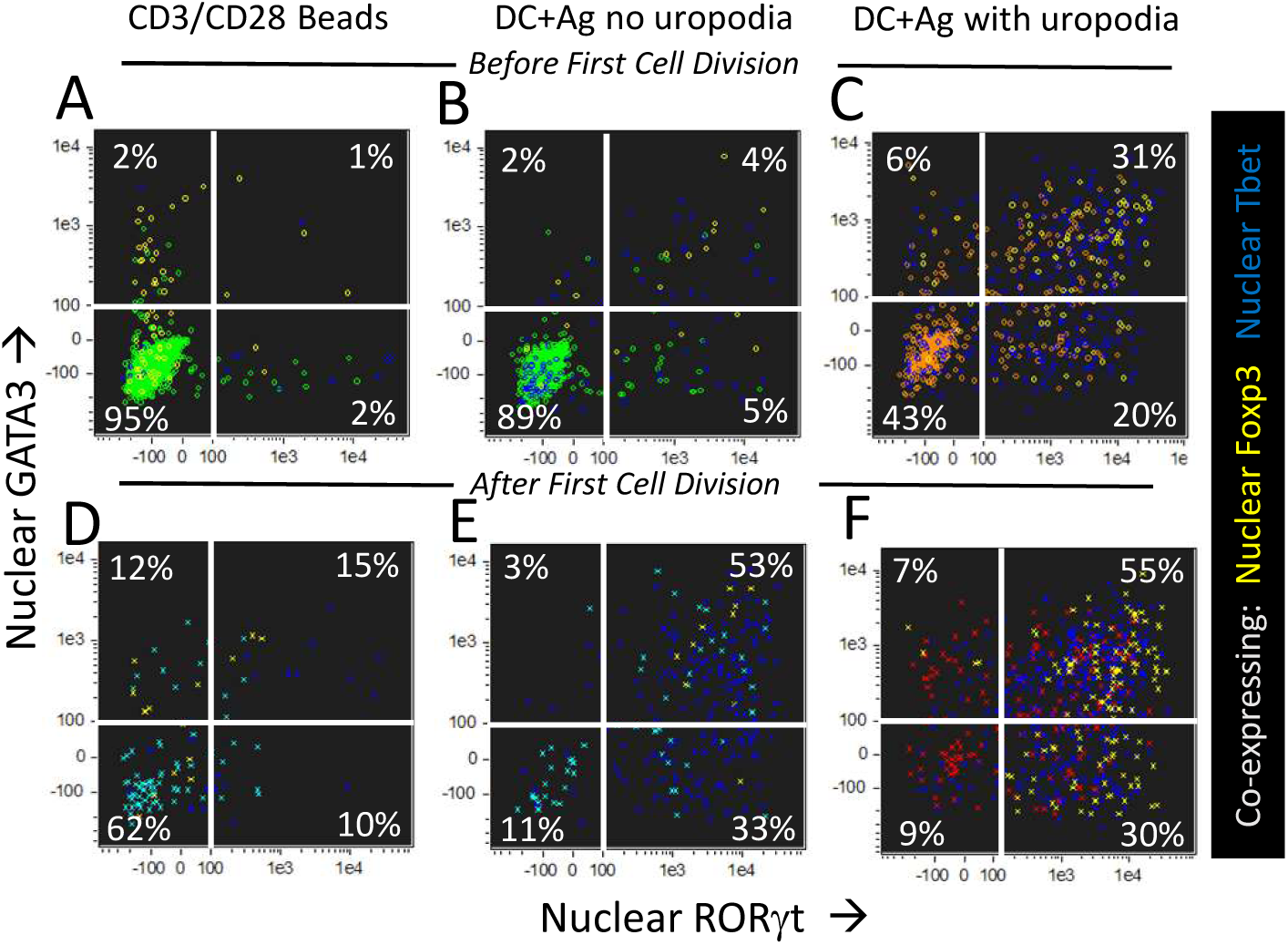
CD4^+^ T cells with uropodia are effector cells co-expressing random combinations of T cell subset transcription factors. **A-F**: CTV labelled female MARKI CD4^+^ T cells were stimulated as indicated with either CD3/CD28 beads (**A, D**) or bmDC + 10nM Dby peptide (**B, C, E, F**), for 2 days with IL-2, TGFβ and ATRA. Cultures were labelled “in situ” with CD4-APC-Cy7, live/dead aqua and Mitotracker DR, fixed and permeabilised, and intracellularly for pS6-Alexa 488, GATA3-PE, RORγt-PE-CF594, Foxp3-PE-Cy7, Tbet-BV605 and 7AAD. 50,000 images were acquired per sample, and gated for live singlet cells in G_0_/G_1_ with (**B, E**: area>10μm^2^ in green) or without (**C, F**: area<10μm^2^ in red) uropodia, and, using CTV dilution, for cells that had not yet divided (**A-C**) or had divided once or more (**D-F**). Dot plots show the intensity of nuclear staining for RORγt versus GATA3, with co-expression of nuclear Foxp3 (yellow) or Tbet (blue) also indicated. One of 5 similar experiments shown (2 with MARKI, 3 with A1RAG).

**Figure 12:**
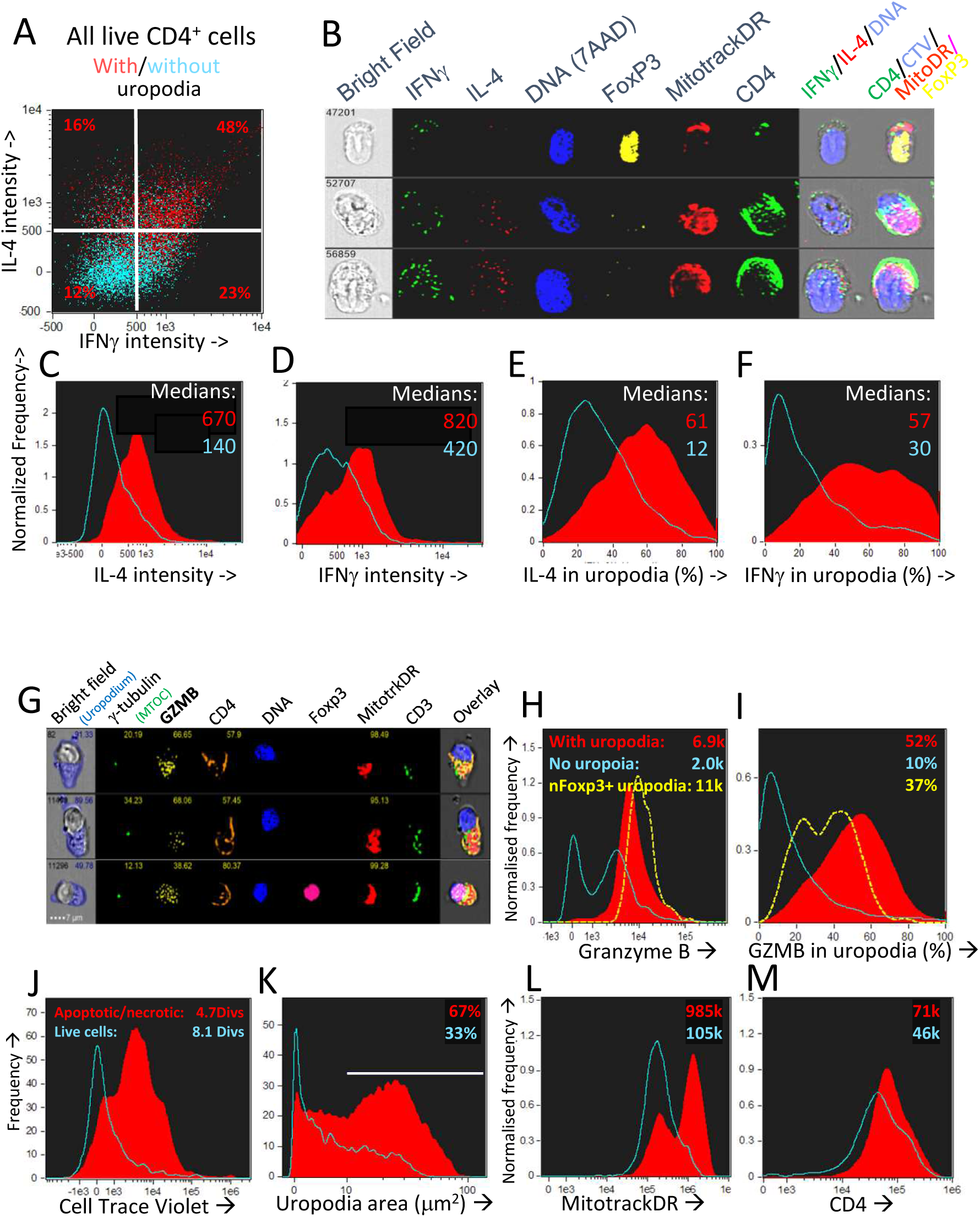
CD4^+^ T cells with uropodia are short-lived effector cells co-expressing random combinations of T cell subset cytokines and granzyme B. **A-F**: CTV labelled female A1RAG CD4^+^ T cells were stimulated in the presence of IL-2, TGFβ and ATRA with bmDC + 100nM Dby peptide for 3 days. Brefeldin was added to the cultures for 2 hours before they were labelled “in situ” with CD4-APC-Cy7, live/dead aqua and Mitotracker DR, fixed and permeabilised, and then intracellularly for IL2-FITC, IFNγ- PE, IL4-PE-CF594, IL17-BV605, Foxp3-PE-Cy7 and 7AAD. Images were gated, as above, for those with (**A**: red dots and **C-F**: red filled histograms) or without (**A**: blue dots and **C-F**: blue histograms) uropodia. Median values of staining intensities for each cytokine are shown in **C-F**, and example images in **B**. One of two similar experiments shown. **G-I**: CTV labelled female A1RAG CD4^+^ T cells were stimulated in the presence of IL-2, TGFβ and ATRA with bmDC + 100nM Dby peptide for 3 days. Cultures were labelled “in situ” with CD4-PE-CF594, CD3-APC-Cy7, CD62L-BV605, Mitotracker DR and live/dead aqua, fixed and permeabilised, and intracellularly stained for γ-tubulin-Alexa488, granzyme B (GZMB– PE) and foxp3-PE-Cy7. Example images are shown in **G**. The intensity of granzyme B staining, with median values indicated (**H**) and the proportion of this staining falling within uropodia (**I**), with median % indicated, for foxp3 negative cells either with (filled red) or without (blue) uropodia, as well as for nuclear foxp3^+^ cells (dashed yellow), are shown (one of two similar experiments). **J-M**: CTV labelled female A1RAG CD4^+^ T cells were stimulated with bmDC + 100nM Dby plus IL2, TGFβ and ATRA for 7 days and labelled “in situ” for Mitotracker DR and live/dead aqua, fixed and permeabilised, and stained for CD4-PE-CF594 and 7AAD. In focus images were gated for singlet cells with a G_0_/G_1_ DNA content. Histograms show the absolute frequencies of live cells (live/dead aqua negative, bright field contrast low: blue histograms) compared to dead/dying cells (apoptotic=bright field contrast high plus necrotic=live/dead aqua positive: filled red) in each plot, comparing cell divisions (**J**), uropodium area (**K**), Mitotracker DR staining (**L**) and CD4 (**M**), with median values indicated. Representative data from many (>10) similar experiments is shown.

### 3.10 Memory T cells make further stochastic cell fate decisions upon TCR restimulation

The cells in the lineage that had never developed uropodia survived and also proliferated until nutrient limitation and mTOR inhibition occurred after day 4 (days 3 and 6 shown: Figure 13A-C). They continued to express high levels of both total and nuclear NFκB even on day 7 (Figure 13D-F), which is thought to be important for the maintenance of memory T cells (49, 57, 58). These putative memory cells continued to survive in a quiescent state until at least day 10 of culture, dependent on IL2 and promoted by TGFβ (Figure 14A-E). By this time almost all the effector cells with uropodia had died (Figure 14D, E). We harvested (on day 6) similar cultures (not previously CTV labelled) and re-stimulated them in fresh medium either with CD3/CD28 beads or DC plus antigen. As expected for memory cells, a lower threshold for activation was evidenced by the fact that CD3/CD28 beads, which were unable to induce uropodia in the primary stimulation (Figure 14I), were sufficient in the secondary stimulation to enable both proliferation and mTOR/pS6 upregulation (not shown) together with uropodium development (Figure 14G). Regardless of the secondary stimulation, we could, once again, observe two distinct populations (either with or without uropodia), suggesting that memory cells make a further “activated/effector” versus “memory/stem” cell fate decision. At the same time, while most of the cells continued to express CD44, a marker for central memory cells (CD62L) was re-expressed on about half of the re-stimulated cells, independent of whether they had developed uropodia or not (Figure 14G, H). This shows that the memory T cells, upon re-stimulation, can apparently make further stochastic, effector/memory-like fate decisions to naïve CD4^+^ T cells. Their T helper and regulatory cell subset transcription factor expression was, however, very different to the random co-expression seen in the primary effector cells, with a much more restricted, singular pattern of either Tbet or Foxp3 in the example shown in Figure 14F.

**Figure 13:**
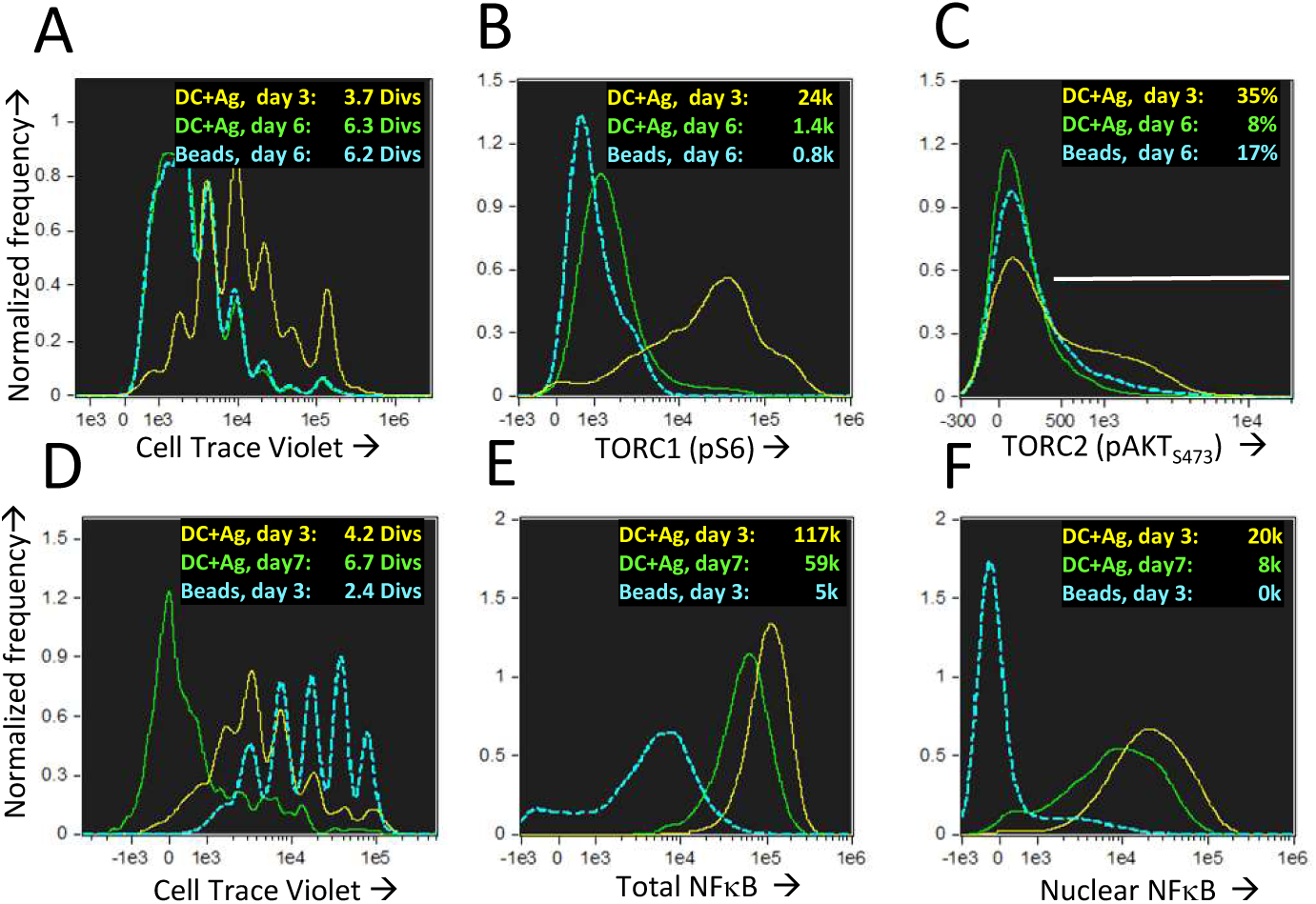
Nuclear NFkB is maintained in long-lived memory cells without uropodia even when they reach quiescence and mTOR activation has ceased. CTV labelled female A1RAG CD4^+^ T cells were stimulated for 3 or 6 days in the presence of IL-2, TGFβ and ATRA with bmDC + 100nM Dby peptide (3 days in yellow, 6 days green) or CD3/CD28 beads (only day 6 shown in blue). “In situ” staining, fixation and image analysis was a previously described, with histograms for CTV (**A**), pS6-Alexa488 (**B**) and pAKTS473 (C) shown. Data from one of many (>10) similar experiments shown. **D-F**: An identical experiment to that above was set up, with DC+Ag stimulation analysed on day 3 (yellow) or day 7 (green) and CD3/CD28 bead stimulation on day3 (blue). Histograms show CTV with mean number of divisions indicated (**D**) and the intensities (with median values shown) of total (**E**) or nuclear (**F**) NFκB (p65)-APC staining. One of two similar experiments.

**Figure 14:**
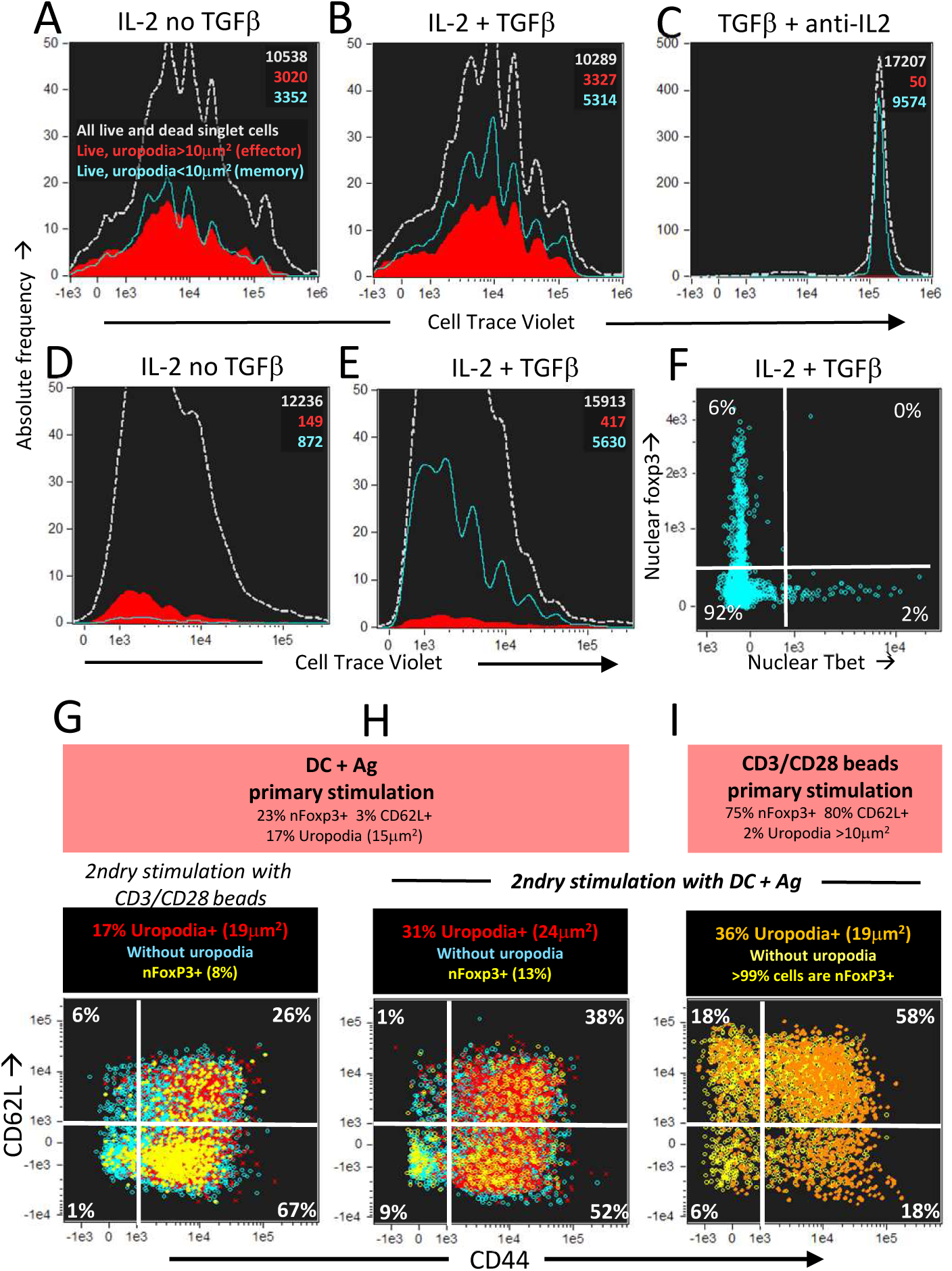
Long-lived memory CD4^+^ T cells, without uropodia, have a lower threshold for re-stimulation, when they make further, independent fate decisions to develop uropodia or re-express CD62L. **A-C**: CTV labelled female A1RAG CD4^+^ T cells were stimulated with bmDC + 100nM Dby for 3 days in the presence of IL2 (50U/ml) without TGFβ (**A**), IL2 (50U/ml) plus TGFβ (2ng/ml) (**B**) or TGFβ plus anti-IL2 (clone S4B6, 50μg/ml) (**C**). Cells were “in situ” labelled with live/dead aqua, fixed and permeabilised, then 7AAD. Histograms show the absolute frequencies of CTV dilution, with grey dashed lines for all images (both live and dead), while filled red (cells with uropodia >10mm2) and blue line (cells without uropodia) histograms are gated for live cells only (live/dead aqua negative, bright field contrast low). Total numbers of cells in each histogram are indicated. One of two similar experiments shown. **D-F**: CTV labelled female MARKI CD4^+^ T cells were stimulated with bmDC + 10nM Dby peptide with IL2 (50U/ml) either with (**E, F**) or without (**D**) TGFβ (2ng/ml) for 10 days. Cells were “in situ” labelled with live/dead aqua and Mitotracker DR (not shown), fixed and permeabilised, then Tbet-Alexa488, foxp3-PE-Cy7, (GATA3-PE, RORγt-PE-CF594, CD4-APC-Cy7, all not shown) and 7AAD. Histograms of absolute frequencies of CTV dilution for total live and dead singlet cells (dashed grey), live cells with (filled red) and without (blue lines) uropodia, together with total numbers of cells are shown (**D, E**). The intensities of Tbet and foxp3 staining within the nucleus of the live cells without uropodia are plotted in **F**. A similar result was also observed using female A1RAG CD4^+^ T cell 10 day cultures (not shown). **G-I**: Female A1RAG CD4^+^ T cells (not CTV labelled) were stimulated in the presence of IL-2, TGFβ and ATRA with bmDC + 100nM Dby peptide (**G, H**) or CD3/CD28 beads (**I**) for 6 days. An aliquot of each sample was analysed as above for uropodia area, CD62L and nuclear foxp3 expression (summarised in orange panels). Cells were harvested, ficoll-hypaque separated, labelled with CTV and Mitotracker DR, then re-stimulated with either CD3/CD28 beads (G) or bmDC+100nM Dby peptide (**H, I**) for 3 days. Cultures were labelled “in situ” with CD4-PE-CF594, CD44-APC-eflour780, CD62L-BV605 and live/dead aqua, fixed and permeabilised, and intracellularly stained for foxp3-PE-Cy7 (Mito-ID-Red, pAKT_S473_-PE, not shown) and Sytox Green for DNA. Plots show the intensity of CD44 and CD62L staining, with % in each quadrant indicated, after gating for live singlet cells with G_0_/G_1_ DNA content, and nfoxp3^−^ cells with uropodia (area >10μm^2^: percentage and median area shown in panel above) in red, without uropodia in blue, and nuclear foxp3^+^ cells (% of all cells shown in panel above) with or without uropodia shown in orange and yellow, respectively. One of 3 similar experiments shown.

## 4. Discussion

### Optimising in vitro culture conditions, fixation, staining and analytical methods

Recent high-profile publications have provided evidence that effector and memory cell fates diverge very early after the activation of both CD4^+^ and CD8^+^ naïve T cells, both in vivo and in vitro (11-14). This divergence is claimed to depend on an initial asymmetric cell division, where one daughter cell preferentially inherits effector cell transcription factors, signalling components and a dependence on glycolysis and anabolic metabolism that drives proliferation and an effector cell fate, while the other daughter remains dependent on oxidative phosphorylation and “defaults” to a memory cell fate. The main weakness in all these publications is the difficulty in directly demonstrating the asymmetric cell divisions, with most data depending on an indirect correlation with high versus low expression of various markers, particularly CD4 or CD8, after the first cell division. Confocal imaging of apparently asymmetric telophases is limited to small numbers of selected images, with a high potential for observer bias, and may be artefactual if cells are not maintained under optimal culture conditions throughout. Imaging flow cytometry data, as we show here, can also be misleading without optimisation of the culture and staining conditions. Most of the published data assumes that an actin bridge between two cells in contact represents cytokinesis, but we have found many such images where such conjugates are clearly between two cells with different CTV dilutions, and so cannot be derived from cell division (data not shown). Disruption of microtubular dynamics, either by using mitotic inhibitors (REF) or even cell harvesting, handling, or non-physiological temperatures (as shown above) can cause asymmetric artefacts or complete loss of classical mitotic figures. Once we had optimised both the culture conditions and the fixation, staining and flow cell imaging analysis, we analysed numerous cell surface markers, transcription factors and signalling molecules across all telophase cells within multiple samples, and never found any evidence of significant asymmetry.

With the increasing realisation in the literature of the importance of T cell metabolism, we needed to better define and control the nutrient and cytokine availability, and therefore moved to a chemically defined medium with only minimal FCS. This meant that any cytokines in the culture were either those we added exogenously or derived from the antigen presenting cells, rather than being a poorly defined “background” source, as may be the case for TGFβ contributed by the higher serum concentrations of traditional culture media. Resulting from this modification it became clear that active TGFβ was important for the “balanced” generation of both effector and memory cells and their survival when stimulated by bmDC plus antigen under these low serum conditions. Active TGFβ addition also had very different effects depending on the context: with CD3/CD28 stimulation it acted as a strong inhibitor of activation and proliferation, while it had the opposite effects with antigen specific stimulation by bmDC.

### Mechanisms of effector versus memory cell fate decisions

One school of thought has been that memory cells emerge as survivors from the expanded pool of effector cells, either due to a switch in the cytokines that support their proliferation or survival, from IL-2 to IL15 (59) or as a result of additional co-stimulatory signals from TNFRSF members (5, 52) such as CD27 and OX40 (CD134). Our experimental system uses medium with only 1% serum, so any cytokines come from the activated T cells themselves, are added exogenously, or come from the antigen presenting dendritic cells. Proliferation was entirely IL2 dependent, whether we stimulated with DC or CD3/CD28 beads, as a neutralising anti-IL2 antibody (S4B6) blocked full activation (cells expressed high CD25 but not CD44) and entry into cell division (Figure 14C). For routine experiments we added sufficient exogenous IL2 such that intrinsic IL2 production would not be a confounding variable to consider. The further addition of IL15 or related cytokines (IL7, IL4) did not change the balance of uropodia expressing effector to memory cells (not shown). We also observed both effector and memory cell populations under Th1 (anti-IL4 plus IFNγ) or Th2 (added IL4 plus anti-IL12) conditions (details not shown, although data from these conditions are shown in Figure 6). The addition of active TGFβ was required for the long-term survival of memory cells after DC + Ag stimulation, as well as any generation of foxp3^+^ Treg cells, especially in the low (1%) serum cultures. A requirement for TGFβ for normal memory cell development in vivo has previously been reported (49, 57, 58). Uropodia development and a bimodal mitochondrial distribution occurred within the first 48 hours after stimulation, and before the initiation of cell proliferation under all the cytokine conditions that we tested.

This seems incompatible with memory cells deriving from a few surviving effector cells. Yet, two recent papers claim exactly that for viral specific memory cells after clinical vaccination (60, 61). How can this be? The first paper makes the common (but we would suggest incorrect) assumption that by labelling proliferating cells in a primary response that these were all effector cells so that if memory cells in a secondary challenge were still labelled they were presumed to derived from them. But we show here that during the primary response the committed memory population are proliferating in parallel to the effector cells. The second paper makes the same assumption and additionally finds that memory cells share some of the epigenetic signature of effector cells, so erroneously claims that memory cells are de-differentiated effectors. But we show that the optimal initial activation of naïve CD4^+^ T cells can induce multiple T cell effector and regulator transcription factors, which would likely leave an epigenetic signature in cells destined to become both effector and memory T cells.

A different view has been that the effector/memory cell fate decision is deterministic and binary, with an extreme example being where an asymmetric cell division generates two daughters, one proliferating to generate effector cells while the other is committed to the memory cell lineage. Our data has similarity to this latter view, except that we did not observe a one-to-one binary fate decision, nor was there any evidence for any asymmetric cell division. Why might this be? We did not use any mitotic inhibitors in our cultures, as these are known to generate artefactual asymmetry during cytokinesis (37). By fixation at 37°C we “froze” any cells in late anaphase/telophase rather than allowing lower temperatures (62), centrifugation or nutrient starvation to disrupt cell structure or cause mitotic spindle collapse which might generate artefactual asymmetry. One limitation was that we were also unable to stain for numb or notch, claimed to be the drivers of asymmetric cell division (63, 64), as a number of commercially (polyclonal) available antibodies gave no cell surface staining above background in our hands, and staining for notch signalling with a monoclonal anti-NICD was also unhelpful (not shown). Notwithstanding this limitation, we found that that the effector lineage was tightly linked to uropodium development before any cell division, and while both lineages were generated across a wide range of conditions, the proportion of effector versus memory cells was regulated in a non-linear manner by signalling through both TCR/CD28, acting primarily through PI3K on mTORC1, but also by GITR (TNFRSF18) signalling via mTORC2/pAKT_S473_ and NF_κ_B. Independently, we found that GITR/mTORC2 signalling and uropodium development were associated with an early (pre-division) induction of random combinations of transcription factors for T helper and regulatory cell subsets, which generate an early cohort of poly-functional effector cells. GITR signalling via mTORC2 was important for uropodium expression and effector cell commitment, but it also led to increased expression of NF__κ__B in all CD4^+^ T cells. NF_κ_B localisation to the nucleus, however, was seen primarily in the long-lived memory cells. This dual role of GITR co-stimulation suggests it might promote the decision process rather than the actual choice of cell fate. The decision to develop uropodia was independent of T cell subset differentiation, and nuclear foxp3-expressing Treg cells were similarly distributed amongst the uropodia positive and negative populations at this early time point, suggesting they too can commit to either an activated/effector or long-lived memory lineage. The Tregs within the effector or memory populations were mTOR/pS6 high or low, respectively, which may explain some of the discrepant reports concerning the status of mTOR in Tregs (65).

### A role for mTOR signalling in cell fate decisions

It has been suggested that mTORC1 and mTORC2 signalling are important for Th2 versus Th1 differentiation, while Tregs develop in the absence of both pathways (66-69). Most of this type of data has been generated using genetic manipulations that completely knock out one or more components of the PI3k/mTOR pathway in T cells, but this fails to take into account that these pathways do not act in a digital fashion but rather they integrate multiple inputs and feedback elements to produce more complex outcomes, as we observed in Figure 9 and (70). For uropodium development, for example, this requires an intermediate level of mTORC1/pS6 but high mTORC2/pAKT_473_, and yet developing Th1, Th2 and Treg seem to be roughly equally distributed amongst the uropodia positive and negative populations on day3, suggesting the mTOR signalling requirements for different T cells subsets may be similar initially. At later time points, nutrient utilisation leads to mTOR inhibition, conditions which may then select for the differential growth and survival of, for example, regulatory rather than inflammatory T helper cell subsets. It is also possible that the 6 discrete mTORC1/mTORC2 signalling populations (Figure 9) are associated with specific T helper and regulatory cell subset differentiation, but we are currently limited by the combinations of reagents and number of fluorescent channels on the ImageStream to test this.

### Autophagy, mitophagy and cell fate

Autophagy is the process by which all cells are able to recycle their cellular components and organelles, including mitochondria (ie. mitophagy), particularly under conditions of stress and nutrient starvation. Autophagy is inherently linked to the mTOR pathway and metabolic status of the cell, and has been implicated in controlling immune cell differentiation (41). Our data suggest that autophagy may be involved in the effector versus memory cell fate decision, as we observed loss of both Cell Trace Violet (which permanently and non-specifically labels intracellular proteins) and Mitotracker DR (which permanently labels intra-mitochondrial proteins) when cells had been pre-labelled with both dyes, and this occurred only after activation (increase in size), before entry into the first cell division, and only with DC plus antigen stimulation. In addition, we could observe staining with a dye that specifically labels late autophagic vesicles at this same time point, at around 48 hours after stimulation with DC + Dby peptide, but not in CD3/CD28 bead stimulated cultures (where we do not see generation of the two cell fates). mTOR inhibition by rapamycin usually induces autophagy in actively growing cells, but in this case we observed reduced staining for autophagy, probably because the primary effect of mTOR inhibition was to reduce and delay the T cell activation required before autophagy could occur. We could not formally prove, however, that the cell fate decision depended specifically on autophagy, as the use of inhibitors such as chloroquin and spautin 1 also blocked T cell activation and proliferation.

### Cell fate decisions after secondary stimulation of memory cells

In our in vitro cultures, nutrient depletion and consequent mTOR inhibition became dominant after 3-5 days of cell proliferation. During this period, effector cells with uropodia died, but could still be observed as cells with reduced DNA staining, increased bright field contrast and live/dead aqua staining, and with a CTV dilution equivalent to an average of 4.2 cell divisions (Figure 12J). Developing memory cells without uropodia, however, continued to proliferate (more than 8 cell divisions by CTV dilution: Figure 12J), became quiescent (as indicated by loss of pS6 staining: Figure 13B), but remained viable and still expressing nuclear NFkB until at least day 10 after stimulation. More importantly, they could harvested, labelled with CTV and re-stimulated, either with CD3/CD28 beads or DC + antigen, to induce a second round of activation and proliferation. Interestingly, around half of the re-activated cells now developed uropodia, regardless of the stimulation (naïve CD4^+^ T cells did not develop uropodia with CD3/CD28 bead stimulation alone), suggesting they could re-capitulate the effector versus fate decision of naïve CD4^+^ T cells to generate “effector memory” and what might be considered a central memory “stem” cell (71). At the same time, around half of the re-activated memory CD4^+^ T cells re-expressed CD62L (which had been lost from naïve T cells after activation) while they remained CD44^+^, compatible with a central memory phenotype. The re-expression of CD62L and the development of uropodia were randomly associated, suggesting these further cell fate decisions were stochastic in nature. Secondary stimulation of memory cells seemed to only give a restricted expression of T cells subset transcription factors, compared to the random co-expression seen on stimulation of naïve T cells. Preliminary experiments outside the main scope of this paper suggest that a cytokine, and mTOR signalling dependent, selection process may operate during memory cell proliferation and development.

### Implications of the model for peripheral Treg development and tolerance

We found a clear bimodal distribution of mitochondrial numbers within CD4^+^ T cells in rejecting mice, while tolerant mice only had a single population with high numbers of mitochondria. It is generally thought that effector cells require glycolysis, while memory and regulatory T cells use oxidative phosphorylatio - this might suggest that the rejecting mice might have both effector and memory T cells in their secondary response, while tolerant mice might lack the glycolytic, low mitochondrial effector population, and have only the regulatory T cells we know are required to maintain tolerance (21). There are, however very few foxp3^+^ Treg cells in the draining lymph nodes of these secondary challenge, tolerant mice as they are likely all within the graft itself (40)), and there are similar numbers of CD44^+^ activated T cells (Figure 4). The in vitro model also shows that it is the effector cells (with uropodia) that correspond to the high mitochondrial containing population, and that the low mitochondria cells that are missing in tolerant mice are more likely to be memory cells. We also showed many years ago that mice tolerant of a skin graft have an increased, rather than decreased, frequency of circulating (splenic) effector Th1 and Th2 cells (22). Taken together, the immune “defect” in tolerant mice would correspond to a lack of memory cells, so that a secondary challenge only elicits a short-lived, primary-like effector cell response in the circulation that can be adequately controlled by the regulatory foxp3^+^ T cells residing within the tolerated skin graft tissue.

### Metabolism and cell fate decisions – cause or effect?

We know that there seems to be a strong link between effector T cells and glycolysis, compared to memory cells and regulatory T cells, which are more dependent on oxidative phosphorylation and fatty acid metabolism. The Advanced RPMI medium we used should provide, at least initially, an excess of metabolic precursors for all the different pathways. It contains a defined content of fatty acids, known to be important for effective memory cell differentiation/survival (23, 24, 72), insulin to ensure effective glycolysis thought to be important for effector cells (15, 73), and a non-labile source of glutamine (important as a carbon source for proliferating T cells). Even so, we found that mTORC1 (pS6) signalling peaked on day 2 (MARKI) or 3 (A1RAG) of culture, after which nutrient limitations increasingly led to mTOR inhibition, apoptosis in the effector cells and a reduction in the rate of memory cell proliferation. We found that uropodia-positive effector cells were higher in both mTORC1 and 2, which would be expected to drive anabolic metabolism, glucose uptake and glycolysis. Conversely, memory cells without uropodia were generally very low/negative for mTORC2 and lower in mTORC1, which would normally be an indication of catabolism, autophagy and OXPHOS (74, 75). It was the effector cells with uropodia, however, which had the higher numbers of mitochondria that stain with Mitotracker DR (that depends on an active electron transport chain), suggesting they may also be active in OXPHOS. We have investigated in some detail the metabolism of these cells committed to either effector or memory cell fates as they developed, but to describe these data in any detail is beyond the scope of this current paper. In particular, it remains an important question whether changes in T cell metabolism that correlate with different differentiation pathways and cell fates are causative or whether they are a result of different metabolic needs, and if the latter, whether the nutrient status of different microenvironments act in a selective manner. We will address these issues in a follow up paper (manuscript in preparation).

This new in vitro model system will enable us to examine how different cytokines, nutrients and other mediators skew responses towards alternative T helper and regulatory subsets as it can distinguish between their relative roles in induction/commitment versus proliferation/selection/survival with important implications for potential therapeutic interventions aimed at manipulating the microenvironment.

## 5. Acknowledgments

We would like to thank the PSB staff for their support with animal care, Nigel Rust for assistance with cell sorting and Annemieke ten Bokum for technical assistance. This work was supported by the MRC UK (Program Grant), the EPA Abraham Trust and the European Research Council (PARIS).

## 6. Author contributions

SPC designed and analysed flow imaging experiments and wrote the manuscript. EA set up and planned in vitro culture and in vivo grafting experiments. DH helped design experiments with mitochondria and writing the manuscript. HW holds grants that funded this research and contributed to experimental design and writing of the manuscript.

